# The *Caulobacter* NtrB-NtrC two-component system bridges nitrogen assimilation and cell development

**DOI:** 10.1101/2023.06.06.543975

**Authors:** Hunter North, Maeve McLaughlin, Aretha Fiebig, Sean Crosson

## Abstract

A suite of molecular sensory systems enables *Caulobacter* to control growth, development, and reproduction in response to levels of essential elements. The bacterial enhancer binding protein (bEBP) NtrC, and its cognate sensor histidine kinase NtrB, are key regulators of nitrogen assimilation in many bacteria, but their roles in *Caulobacter* metabolism and development are not well defined. Notably, *Caulobacter* NtrC is an unconventional bEBP that lacks the σ^54^- interacting loop commonly known as the GAFTGA motif. Here we show that deletion of *C. crescentus ntrC* slows cell growth in complex medium, and that *ntrB* and *ntrC* are essential when ammonium is the sole nitrogen source due to their requirement for glutamine synthetase (*glnA*) expression. Random transposition of a conserved IS3-family mobile genetic element frequently rescued the growth defect of *ntrC* mutant strains by restoring transcription of the *glnBA* operon, revealing a possible role for IS3 transposition in shaping the evolution of *Caulobacter* populations during nutrient limitation. We further identified dozens of direct NtrC binding sites on the *C. crescentus* chromosome, with a large fraction located near genes involved in polysaccharide biosynthesis. The majority of binding sites align with those of the essential nucleoid associated protein, GapR, or the cell cycle regulator, MucR1. NtrC is therefore predicted to directly impact the regulation of cell cycle and cell development. Indeed, loss of NtrC function led to elongated polar stalks and elevated synthesis of cell envelope polysaccharides. This study establishes regulatory connections between NtrC, nitrogen metabolism, polar morphogenesis, and envelope polysaccharide synthesis in *Caulobacter*.

**Importance:** Bacteria balance cellular processes with the availability of nutrients in their environment. The NtrB-NtrC two-component signaling system is responsible for controlling nitrogen assimilation in many bacteria. We have characterized the effect of *ntrB* and *ntrC* deletion on *Caulobacter* growth and development and uncovered a role for spontaneous IS element transposition in the rescue of transcriptional and nutritional deficiencies caused by *ntrC* mutation. We further defined the regulon of *Caulobacter* NtrC, a bacterial enhancer binding protein, and demonstrate that it shares specific binding sites with essential proteins involved in cell cycle regulation and chromosome organization. Our work provides a comprehensive view of transcriptional regulation mediated by a distinctive NtrC protein, establishing its connection to nitrogen assimilation and developmental processes in *Caulobacter*.

## Introduction

Nitrogen exists in a multitude of forms in the environment, and bacteria have a variety of molecular mechanisms to assimilate this essential element. Accordingly, bacterial cells commonly express sensory transduction proteins that detect environmental nitrogen and regulate the transcription of genes that function in nitrogen assimilation. The conserved NtrB-NtrC two-component system (TCS) is among the most highly studied of these regulatory systems. The NtrB-NtrC TCS has been broadly investigated, particularly in Enterobacteriaceae where it is well established that the NtrB sensor histidine kinase controls phosphorylation state of the DNA-binding response regulator, NtrC, in response to intracellular nitrogen and carbon status (1-4). Phospho-NtrC (NtrC∼P) activates transcription of multiple genes involved in inorganic nitrogen assimilation and adjacent physiologic processes.

The preferred inorganic nitrogen source for many bacteria is ammonium (NH_4_^+^) (5), and NtrC∼P commonly activates transcription of glutamine synthetase (*glnA*) (6), which functions to directly assimilate NH_4_^+^ in the process of glutamine synthesis. In the freshwater-and soil-dwelling bacterium, *Caulobacter crescentus* (hereafter *Caulobacter*) (7), glutamine levels *per se* are a key indicator of intracellular nitrogen status and impact cell differentiation and cell cycle progression via the nitrogen-related phosphotransferase (PTS^Ntr^) system (8). The deletion of *ntrC* results in a nitrogen deprivation response in *Caulobacter* (8) and it is expected that this is due, at least in part, to reduced *glnA* transcription. However, NtrC belongs to a broadly conserved class of transcriptional regulators known as bacterial enhancer binding proteins (bEBPs) that can function as global regulators of gene expression (9), so NtrC is predicted to regulate expression of more than just *glnA* in *Caulobacter*. Indeed, ChIP-seq and transcriptomic studies in *Escherichia coli* demonstrated that NtrC binds dozens of sites on the chromosome (10, 11) and affects transcription of »2% of the genome (12). Given the importance of cellular nitrogen status as a cell cycle and developmental regulatory cue in *Caulobacter*, we sought to define the NtrC regulon and to assess the role of the NtrB-NtrC TCS in the regulation of cell development and physiology.

*Caulobacter* NtrC is a standard (Group 1) bEBP and thus possesses, a) a receiver (REC) domain, b) an ATPase associated with cellular activity (AAA+) domain, and c) a DNA-binding/helix-turn-helix (HTH) domain (9) (Fig S1). Phosphorylation of a conserved aspartate residue in the REC domain regulates oligomerization of Group 1 bEBPs as transcription factors, which activate transcription from s^54^-dependent promoters through a mechanism that requires ATP hydrolysis by the AAA+ domain (9). However, there are limited examples of NtrC proteins that lack a critical series of amino acids in the AAA+ loop 1 (L1) domain known as the GAFTGA motif, which is necessary for interaction with the s^54^ N-terminal regulatory domain (13). Some bEBP proteins lacking the L1 GAFTGA motif are reported to regulate transcription from s^70^-dependent promoters through a mechanism that does not require ATP hydrolysis (14-16). Species in the genus *Caulobacter* harbor a distinct eight residue deletion in AAA+ L1 of NtrC that encompasses the GAFTGA motif (Fig S1). This observation raised the question of whether *Caulobacter* NtrC has functional/regulatory properties that are distinct from orthologs of other genera.

We conducted a molecular genetic analysis of the *Caulobacter* NtrB-NtrC TCS. A main objective of this study was to determine the functional roles of *ntrB* and *ntrC* during growth in media containing inorganic and organic nitrogen sources. Using transcriptomic and ChIP-seq approaches, we defined the NtrC regulon, revealing its dual function as both an activator and a repressor. Our ChIP-seq analysis identified dozens of NtrC binding sites across the *Caulobacter* chromosome, many of which directly overlap with binding sites for the essential nucleoid-associated protein, GapR (17, 18), and the cell cycle regulator, MucR1 (19). Deletion of *ntrC* led to slow growth in complex medium and an inability to grow when NH_4_^+^ was the sole nitrogen source, due to a lack of *glnBA* transcription. Random transposition of a conserved *Caulobacter* IS3-family mobile genetic element into the promoter of the *glnBA* operon was a frequent and facile route to rescue the growth defect of *ntrC* mutants; IS3 transposition effectively rescued *glnBA* transcription, enabling growth of the Δ*ntrC* strain. *Caulobacter* is a prosthecate bacterium that elaborates a thin stalk structure from its envelope at one cell pole, and we further discovered that loss of *ntrC* resulted in hyper-elongated stalks and a hyper-mucoid phenotype. These phenotypes were complemented by either glutamine supplementation to the medium or by ectopic *glnBA* expression. Our study provides a genome-scale view of transcriptional regulation by a NtrC protein with distinct structural features and defines a regulatory link between NtrC and nitrogen assimilation, polar morphogenesis, and envelope polysaccharide synthesis in *Caulobacter*.

## Results

### The nitrogen assimilation defect of *Caulobacter* D*ntrC* is not genetically complemented by *Escherichia coli* or *Rhodobacter capsulatus ntrC*

Given the well-established role for the NtrB-NtrC TCS in inorganic nitrogen assimilation (20), we predicted that a *Caulobacter* mutant harboring an in-frame deletion of *ntrC* (D*ntrC*) would exhibit growth defects in a defined medium with NH_4_^+^ as the sole nitrogen source (M2 minimal salts with glucose; M2G). As expected, the D*ntrC* mutant failed to grow in M2G and this growth defect was genetically complemented by restoring *ntrC* at an ectopic locus (Fig 1A). The sole predicted route of NH_4_^+^ assimilation in *Caulobacter* is via glutamine synthetase (8), and we, therefore, predicted that replacement of NH_4_^+^ with glutamine as the nitrogen source would restore growth of D*ntrC* in M2G. As expected, replacement of NH_4_^+^ with molar-equivalent levels of glutamine (9.3 mM final concentration) restored Δ*ntrC* growth in M2G (Fig 1A). We conclude that *ntrC* is required for NH_4_^+^ assimilation in a defined me-dium.

**Figure 1.**
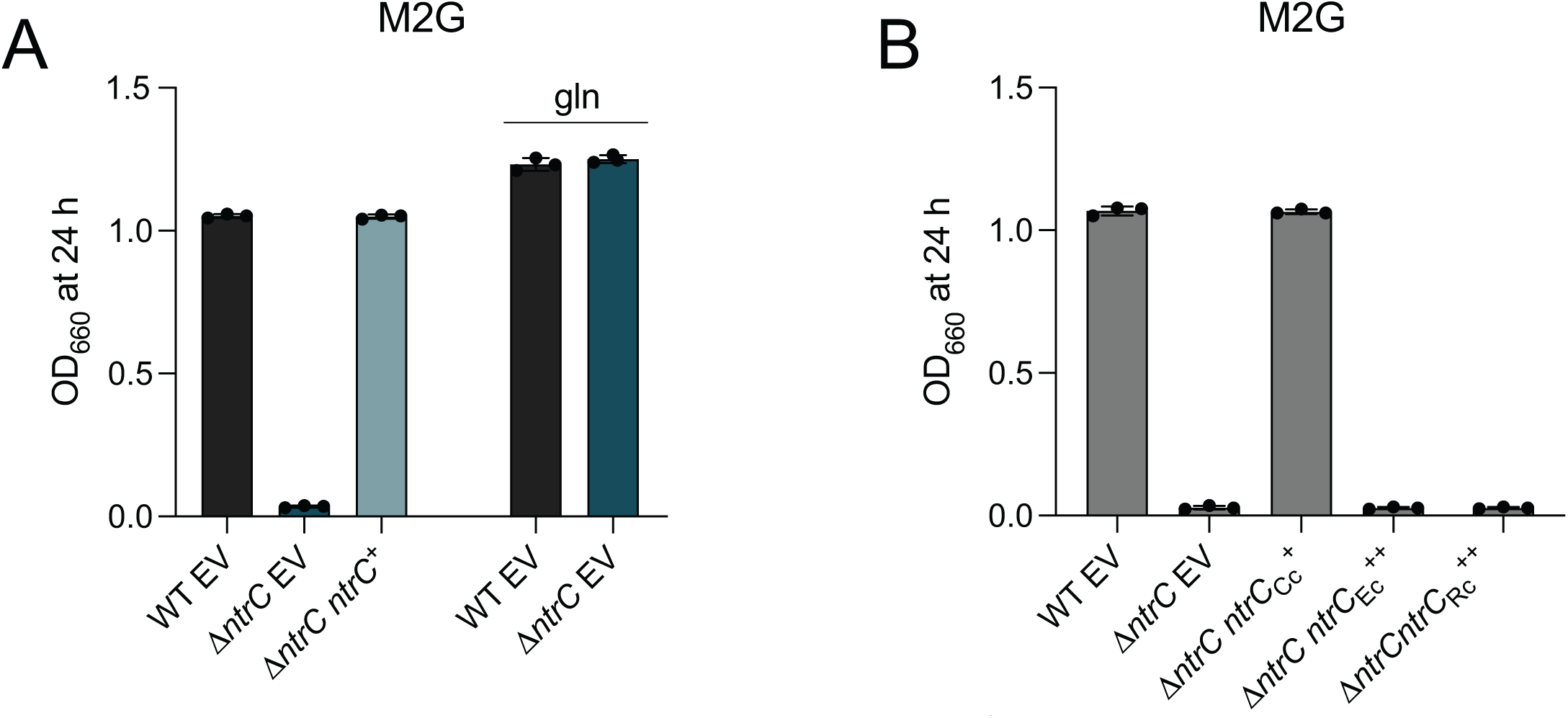
***ntrC* is required for growth in defined medium in which NH_4_^+^ is the sole nitrogen source**. (A) Terminal culture densities of WT, Δ*ntrC*, and Δ*ntrC* carrying a complementing copy (*ntrC*^+^) or empty vector control (EV). Culture growth was measured spectrophotometrically at 660 nm (OD_660_) after 24 hours (h) of growth in M2G or M2G in which NH_4_^+^ was replaced with molar-equivalent (9.3 mM final concentration) glutamine (gln). Data represent mean ± standard deviation of three replicates. (B) Terminal densities of WT and Δ*ntrC* containing empty vector (EV) or expressing *Caulobacter ntrC* from its native promoter (*ntrC*^+^) or *E. coli ntrC* or *R. capsulatus ntrC* expressed from P*_xyl_* (*ntrC_Ec_*^++^ or *ntrC*_Rc_^++^). Culture growth was measured spectrophotometrically at OD_660_ after 24 h of growth in M2G supplemented with 0.15% xylose. Data represent mean ± standard deviation of three independent replicates.

The functional conservation of *ntrC* between phylogenetically proximal (21-23) and distal (24) species has been demonstrated by heterologous genetic complementation. *Caulobacter* NtrC shares 40% sequence identity with the highly studied *Escherichia coli* NtrC (Fig S1), but expression of *E. coli ntrC* from a xylose-inducible promoter did not restore growth of *Caulobacter ΔntrC* in M2G (Fig 1B) even though *E. coli* NtrC was stably produced in *Caulobacter* (Fig S2A). Inspection of NtrC primary sequences revealed that the AAA+ domain from *Caulobacter* species lacks the conserved GAFTGA motif (Fig S1), which is important for the promoter remodeling activity of the AAA+ domain and for coupling promoter conformation information to s^54^-RNAP (13). *Rhodobacter capsulatus*, like *Caulobacter*, is in the class Alphaproteobacteria. NtrC from this species and others in the order Rhodobacterales also harbor a deletion of the L1 loop containing the GAFTGA motif (Fig S1); *R. capsulatus* NtrC is reported to activate gene expression through s^70^ rather than s^54^ (15). Expression of *R. capsulatus ntrC* from a xylose-inducible promoter also failed to restore growth of *Caulobacter ΔntrC* in M2G (Fig 1B), though the protein was stably produced (Fig S2A). The L1 deletion surrounding the GAFTGA motif in *R. capsulatus* NtrC differs – and is larger than – the deletion in *Caulobacter* NtrC (Fig S1). These results provide evidence that *Caulobacter* NtrC has distinct structural and functional features, which merit further investigation.

### Mutation of *ntrB* and *ntrC* has disparate effects on growth in defined versus complex medium

We demonstrated that *ntrC* is essential in M2G defined medium. Glutamine levels in peptone yeast extract (PYE) – a complex medium – are reported to be low (8), and we have confirmed a previous report by Ronneau and colleagues (8) that Δ*ntrC* has a growth defect in PYE that is complemented by expression of *ntrC* from an ectopic locus (Fig 2A) or by addition of glutamine to the medium (Fig 2B). We predicted that deletion of the gene encoding NtrB, the sensor kinase that phosphorylates NtrC *in vitro* (25), would result in similar defects as deletion of *ntrC*. We created an in-frame deletion of *ntrB* (Δ*ntrB*) and observed no effect on growth rate in complex medium relative to wild type (WT) (Fig 2A).

Given this result, we explored the possibility that phosphorylation is not required for NtrC-dependent growth regulation in complex medium. To assess the functional role of NtrC phosphorylation, we mutated the conserved aspartyl phosphorylation site in the receiver domain of NtrC to either alanine (*ntrC*^D56A^), which cannot be phosphorylated, or glutamic acid (*ntrC*^D56E^), which functions as a “phosphomimetic” mutation in some cases (26). Like Δ*ntrB*, the growth rates of *ntrC*^D56A^ and *ntrC*^D56E^ strains were indistinguishable from WT in complex medium, though *ntrC*^D56A^ cultures had reduced terminal density (Fig 2C) that was complemented by glutamine supplementation to the medium (Fig 2D). Both NtrC point mutants were stably produced in *Caulobacter* as determined by western blot (Fig S2B). In fact, steady-state levels of NtrC were elevated in Δ*ntrB* and *ntrC*^D56A^ compared to WT and *ntrC*^D56E^ (Fig S2B), indicating that these proteins are either more stable, more highly expressed, or both. We further investigated growth of these mutants in M2G defined medium. The Δ*ntrB* and *ntrC*^D56A^ strains failed to grow in M2G, while *ntrC*^D56E^ grew like WT (Fig 2G). Like Δ*ntrC*, the growth defect of Δ*ntrB* and *ntrC*^D56A^ in M2G was rescued by replacing NH_4_^+^ with molar-equivalent glutamine (Fig 2H). We conclude that while NtrC phosphorylation does not greatly impact growth in complex medium, it is essential for growth when NH_4_^+^ is the sole nitrogen source.

**Figure 2.**
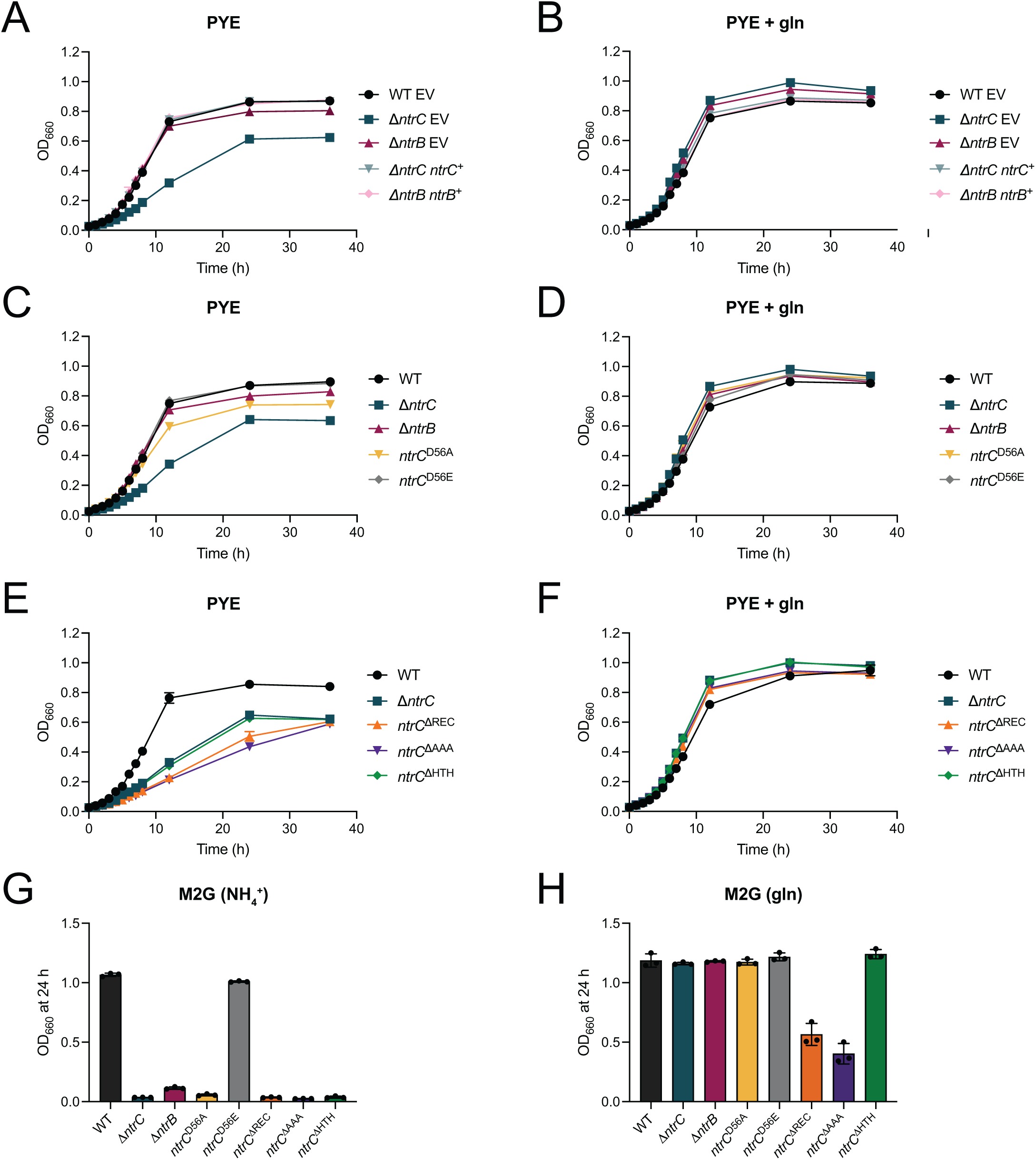
Mutants of the *ntrB-ntrC* system have disparate effects on growth in defined versus complex medium. (A) Growth of WT, Δ*ntrC*, and Δ*ntrB* possessing empty vector (EV) or a genetic complementation vector (^+^) in which indicated genes were expressed from their native promoters (ectopically integrated at the *xylX* locus); growth was measured spectrophotometrically at 660 nm (OD_660_) in PYE complex medium without and (B) with supplemented 9.3 mM glutamine (gln). (C) Growth curves of WT, Δ*ntrC*, Δ*ntrB*, *ntrC*^D56A^, and *ntrC*^D56E^ in PYE and (D) PYE supplemented with 9.3 mM gln. (E) Growth curves of WT, Δ*ntrC*, *ntrC*^ΔREC^ (residues deleted: 17-125), *ntrC*^ΔAAA^ (residues deleted: 159- 363), *ntrC*^ΔHTH^ (residues deleted: 423-462) in PYE and (F) PYE supplemented with 9.3 mM gln. Plotted points for A-F represent average OD_660_ ± standard deviation of three independent replicates. (G) Terminal OD_660_ of WT, Δ*ntrC*, Δ*ntrB*, *ntrC*^D56A^, *ntrC*^D56E^, *ntrC*^ΔREC^, *ntrC*^ΔAAA^, and *ntrC*^ΔHTH^ after 24 hours (h) of growth in M2G defined medium and (H) M2G in which NH_4_^+^ was replaced with molar-equivalent (9.3 mM) gln. Data represent mean ± standard deviation of three inde-pendent replicates.

To extend our structure-function analysis of *Caulobacter ntrC*, we engineered mutant strains harboring *ntrC* alleles in which either the receiver domain (*ntrC*^DREC^; residues 17-125), the s^54^-activating/AAA ATPase domain (*ntrC*^DAAA^; residues 159-363), or the helix-turn-helix DNA-binding domain (*ntrC*^DHTH^; residues 423-462) were removed. Growth of all three mutants (*ntrC*^DREC^, *ntrC*^DAAA^ and *ntrC*^DHTH^) was slower than WT in PYE complex medium, though the growth defects of *ntrC*^DREC^ and *ntrC*^DAAA^ were more extreme than *ntrC*^DHTH^ and D*ntrC* (Fig 2E). The growth defects of all domain mutants in PYE were complemented by glutamine supplementation to the medium (Fig 2F). Each of these domain truncation alleles was stably expressed in *Caulobacter* (Fig S2C). Again, steady-state levels of NtrC^DHTH^, NtrC^DREC^, and NtrC^DAAA^ were elevated, indicating that these mutant proteins are either more stable, more highly expressed, or both. All *ntrC* domain mutants failed to grow in M2G defined medium (Fig 2G). Replacement of NH_4_^+^ with molar-equivalent glutamine in M2G fully rescued the culture yield (i.e. terminal density) of *ntrC*^ΔHTH^, although, yields of *ntrC*^ΔREC^ and *ntrC*^ΔAAA^ were only partially rescued (Fig 2H). Altogether, these results provide evidence that each of the NtrC domains is required for proper NH_4_^+^ assimilation, though the culture yield defects of NtrC^ΔREC^ and NtrC^ΔAAA^ are not solely linked to nitrogen availability.

Having shown that the AAA+ domain of NtrC is required for growth in defined medium, we next investigated the role of ATP binding and ATP hydrolysis by this domain in NH_4_^+^ assimilation. To probe the impact of ATP binding on NtrC function, we mutated the conserved lysine (K178) in the Walker A motif of AAA+, which is necessary for ATP binding in bEBPs (27) (Fig S1). To evaluate ATP hydrolysis, we mutated the conserved aspartate residue (D235) within the Walker B motif of AAA+ (Fig S1). This residue is vital for ATP hydrolysis but not for ATP binding (27, 28). Strains solely expressing either the *ntrC*^K178A^ or *ntrC*^D235A^ alleles did not grow in M2G defined medium (Fig S3), though these mutant proteins were stably expressed in *Caulobacter* (Fig S2D). As observed in other null NtrC mutants, steady state levels of NtrC^K178A^ and NtrC^D235A^ were increased, suggesting that these proteins are either more stable, more highly expressed, or both. These results provide evidence that conserved residues in the NtrC AAA+ domain known to impact ATP binding and ATP hydrolysis are required for NH_4_^+^ assimilation in defined medium.

### IS3 rescue of *glnBA* transcription restores growth of **Δ*ntrC*.**

During our investigation of Δ*ntrC*, we noticed occasional instances of robust bacterial growth in M2G defined medium, indicating the possibility that spontaneous mutation(s) could bypass the growth defect of Δ*ntrC*. Indeed, in four independent cases in different *ntrC* mutant backgrounds, we isolated suppressor mutants that exhibited growth in M2G (Table S1 and Fig 3A). Whole genome sequencing revealed that in three of these strains, an IS3- family (IS511/ISCc3) insertion element had integrated into the promoter of the *glnBA* operon. In the Δ*ntrC* parent strain, an IS3 family insertion element inserted 8 bp upstream of *glnB* (Δ*ntrC* P*_glnBA_*::IS3) (Fig 3B); this insertion was accompanied by a large deletion of sequence in the adjacent operon (*CCNA_02043*-*02045*). We also identified two independent IS3-family (IS511/ISCc3) insertions upstream of *glnBA* (16 bp and 51 bp upstream of *glnB*) that rescued the growth defect of *ntrC*^ΔHTH^ mutants. In di-verse bacteria, NtrC∼P is known to activate transcription of *glnA* (6), which encodes glutamine synthetase. This enzyme directly assimilates NH_4_^+^ by synthesizing glutamine from NH_4_^+^ and glutamate. *glnB* encodes a conserved PII protein that regulates GlnA (6). We observed a fourth growth rescue mutation in *ntrC*^D56A^, where a non-synonymous intragenic transversion resulting in a N94Y mutation rescued growth of the non-phosphorylatable NtrC^D56A^ mutant (Table S1).

**Figure 3.**
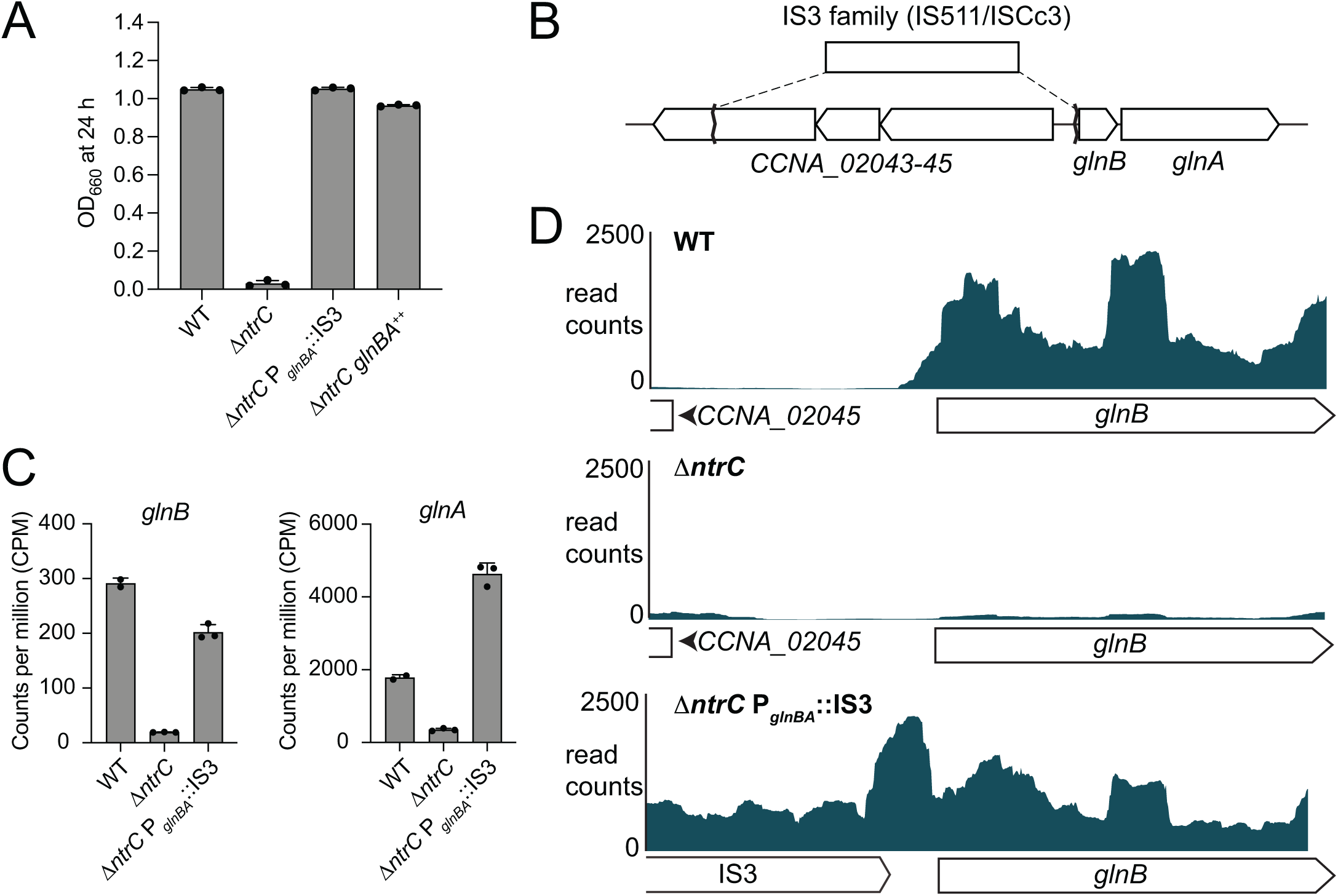
Spontaneous transposition of an IS3-family insertion element restores *glnBA* expression in Δ*ntrC* and rescues the Δ*ntrC* growth defect. (A) Terminal optical density (OD_660_) of WT, Δ*ntrC*, a spontaneous suppressor of Δ*ntrC* (Δ*ntrC* P*_glnBA_*::IS3), and Δ*ntrC* expressing *glnBA* from an inducible promoter (Δ*ntrC glnBA*^++^) grown for 24 hours (h) in M2G defined medium. (B) Site of the spontaneous lesion upstream of *glnBA* in the Δ*ntrC* suppressor strain as determined by whole-genome sequencing. The insertion sequence (IS) element in inserted such that the 3’ end of the transposase (matching the 3’ end of the transposases *CCNA_00660* and *CCNA_02814*) is positioned at nucleotide 2192508, which is 8 nucleotides upstream of the *glnB* start codon. In addition, a 3285 bp deletion eliminated most of *CCNA_02043-45* operon. (C) RNAseq counts per million (CPM) of *glnB* (left) and *glnA* (right) transcripts in WT, Δ*ntrC*, and Δ*ntrC* P*_glnBA_*::IS3 exponential phase cells grown in PYE complex medium. (D) Aligned RNAseq read counts (blue) corresponding to the 5’ end of the *glnBA* operon from WT, Δ*ntrC*, and Δ*ntrC* P*_glnBA_*::IS3 cells. Annotated regions are diagramed below the x-axis for each strain.

To determine the transcriptional consequences of IS3 insertion at P*_glnBA_*, we assessed global transcript levels in WT, Δ*ntrC*, and the Δ*ntrC* P*_glnBA_*::IS3 suppressor strain (sup 1; Table S1). As expected, the Δ*ntrC* strain had negligible *glnBA* transcripts compared to WT (Fig 3C and Table S2). However, *glnB* and *glnA* transcription was restored in Δ*ntrC* P*_glnBA_*::IS3 (Fig 3C and Table S2). Mapped reads demonstrated transcription originating from the IS3 element that extended into *glnBA* (Fig 3D). This provides evidence that sequences within the IS511-ISCc3 mobile element promote transcription of *glnBA* independent of NtrC, thereby enabling growth of the *Caulobacter* Δ*ntrC* mutant in M2G. To test if *glnBA* transcription alone is sufficient to restore Δ*ntrC* growth, we expressed *glnBA* from a xylose-inducible promoter (Δ*ntrC glnBA*^++^). We ob-served similar growth restoration in M2G in this strain (Fig 3A). These findings demonstrate that the inability of Δ*ntrC* to grow when NH_4_^+^ is the sole nitrogen source is from the lack of *glnBA* transcription, and that this transcriptional (and growth) defect can be rescued by insertion of mobile DNA elements into the *glnBA* promoter.

Considering that strains with mutations affecting NtrC phosphorylation (e.g., Δ*ntrB*, *ntrC*^D56A^) do not grow in M2G (Fig 2G), we examined the effect of *ntrB* and *ntrC* mutations on *glnBA* expression using a fluorescent P*_glnBA_* tran-scriptional reporter (P*_glnBA_*-*mNeonGreen*). Reporter activity was significantly reduced in Δ*ntrB*, *ntrC*^D56A^, and *ntrC*^D56E^ when cultivated in complex medium, although *ntrC*^D56E^ had higher P*_glnBA_*-*mNeonGreen* transcription than *ntrC*^D56A^ (Fig S4). These results provide evidence that an intact phosphorylation site in the NtrC receiver domain (D56) is important for the activation of *glnBA* transcription by NtrC. The lack of P*_glnBA_* activity in Δ*ntrB* supports a model in which NtrB functions as the NtrC kinase *in vivo*.

### Defining the *Caulobacter* NtrC regulon

NtrC belongs to a class of proteins known as bEBPs, which often function as global regulators of transcription in bacteria. We sought to comprehensively define the NtrC regulon in *Caulobacter*. To this end, we used RNA sequencing (RNA-seq) and chromatin immunoprecipitation sequencing (ChIP-seq) approaches. Deletion of *ntrC* significantly changed transcript levels for nearly one-quarter of genes in the *Caulobacter* genome relative to WT (RNA-seq; *P* < 10^-4^) when strains were cultivated in PYE complex medium (Fig 4A and Table S2). To distinguish genes directly regulated by NtrC from indirectly regulated genes, we performed ChIP-seq using a 3xFLAG-tagged NtrC fusion. This experiment identified 51 significantly enriched peaks (Fig 4D and Table S3), which represent direct NtrC binding sites. From these peaks, we identified a common DNA sequence motif (Fig 5A) that is significantly related to the multifunctional DNA-binding protein Fis of *E. coli*, and with the NtrC motif of *E. coli*, though there are features that clearly distinguish the *Caulobacter* NtrC motif from *E. coli* NtrC (Fig S5).

**Figure 4.**
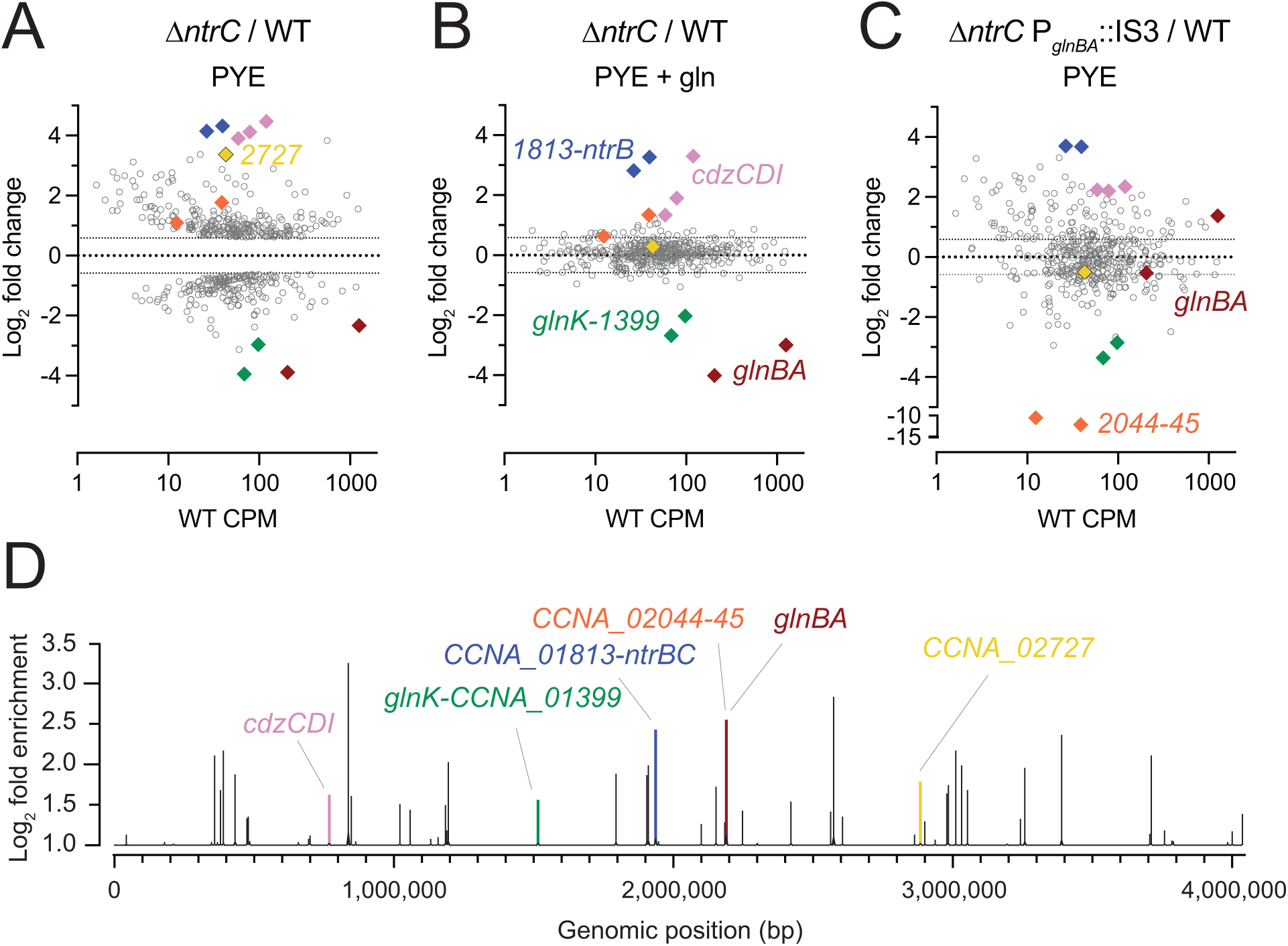
NtrC globally regulates gene expression in *Caulobacter*. In PYE complex medium, 473 genes exhibit differential transcript abundance in the Δ*ntrC* mutant compared to WT based on the following criteria: fold change > 1.5, FDR *P* < 0.000001 and maximum group mean RPKM > 10. (A-C) Log_2_ fold change in abundance for these 473 transcripts for the following comparisons: (A) Δ*ntrC* vs. WT cultures grown in PYE, (B) Δ*ntrC* vs. WT cultures grown in PYE supplemented with 9.3 mM glutamine (gln). Differentially regulated genes in the Δ*ntrC* mutant are largely restored to WT-like levels upon supplementation with glutamine; exceptions are highlighted in colored diamonds. (C) Δ*ntrC* P*_glnBA_*::IS3 vs. WT cultures grown in PYE, where each symbol represents a gene. The x-axis represents WT transcript abundance in PYE (counts per million; CPM) for each gene. (D) NtrC ChIP-seq peaks (q-value < 0.05, area under the curve (AUC) > 20) across the *Caulobacter* genome plotted as log_2_ fold enrichment in read counts compared to the input control. Peaks highlighted in color are in the promoter of genes highlighted in (A-C), or, in the case of *cdzCDI*, overlapping the coding region. Colors correspond to the following genes: pink, *cdzCDI*; green, *glnK*-*CCNA_01399*; blue, *CCNA_01813*-*ntrB*; orange, *CCNA_02044*-*45*; red, *glnBA*; yellow, *CCNA_02727*.

**Figure 5.**
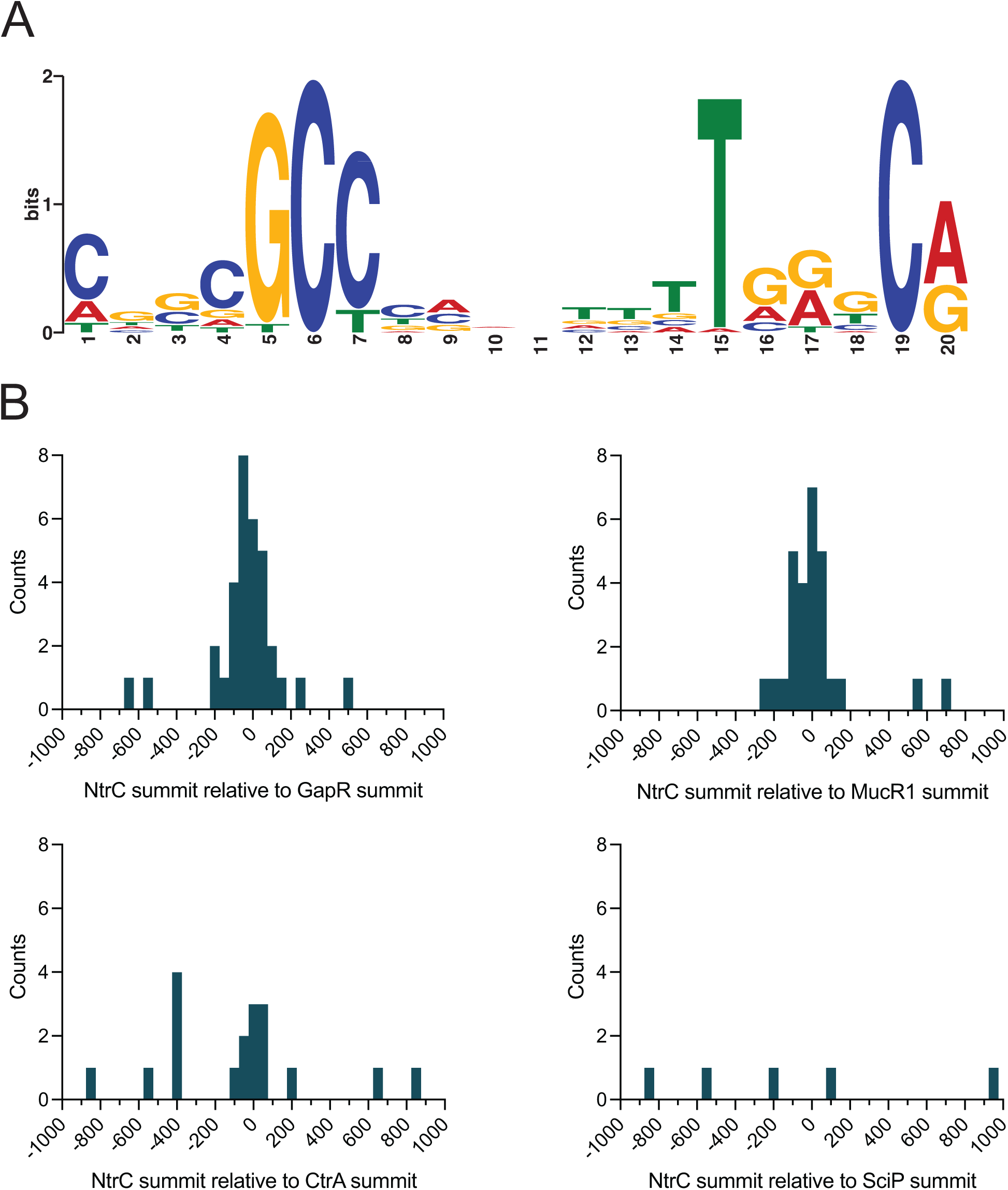
NtrC binding sites are often co-located with the binding sites of select chromosome structuring proteins and cell cycle regulators. (A) DNA motif enriched in NtrC ChIP-seq peaks, as identified by MEME (79). (B) Distribution of the relative position of NtrC ChIP-seq summits to the nearest GapR, MucR1, CtrA, or SciP summit as calculated by ChIPpeakAnno (78). NtrC summits >1,000 bp away from the nearest cell cycle regulator summits were excluded from the plots. Frequency distributions were plotted as histograms with 50 bp bins.

As expected, the data indicate that NtrC directly activates *glnBA*: a major NtrC peak was identified in the *glnBA* promoter region (Fig 4D and Table S3). NtrC also directly binds the promoter region of the *glnK*-*CCNA_01399* op-eron (Fig 4A and Table S3). *glnK* encodes a PII protein homologous to GlnB, which has been shown to similarly regulate GlnA in bacteria (6, 29), while *CCNA_01399* is an annotated as an AmtB-family NH_4_^+^ transporter. Transcript levels of *glnK* and *CCNA_01399* are decreased 8- 15 fold (*P* < 10^-61^), respectively, in Δ*ntrC* relative to WT (Fig 4A and Table S2); we conclude that NtrC directly activates transcription of these genes. We further observed an NtrC peak in the promoter regions of two genes in the nitrate assimilation locus, which is transcriptionally activated by nitrate (30) and functions to reduce nitrate to ammonium. Specifically, NtrC peaks are present at the 5’ end of the nitrate response regulator NasT and in the promoter region of the MFS superfamily nitrate/nitrite transporter NarK (Table S3). RNA-seq measurements were conducted in the absence of nitrate so, as expected, we did not observe differential transcription of this locus. Transcription of genes residing in the same operon as *ntrC*, including *ntrB* and a predicted tRNA-dihydrouridine synthase (*CCNA_01813*), are increased in Δ*ntrC* by approximately 20-fold (Fig 4A and Table S2). NtrC directly binds the promoter of its operon (Fig 4D and Table S3), providing evidence that it functions as an autorepressor. This is consistent with our western blot data showing that *ntrB-ntrC* loss-of-function mutants (e.g., Δ*ntrB*, *ntrC*^D56A^, *ntrC*^ΔREC^, *ntrC*^ΔAAA^, *ntrC*^ΔHTH^, *ntrC*^K178A^, *ntrC*^D235A^) have increased levels of NtrC protein (Fig S2B-D), indicating loss of autorepression at this genetic locus.

We have further identified genes in our datasets that are not known to be directly involved in nitrogen assimilation. In fact, 9 of the 51 NtrC binding sites are located within a mobile genetic element (MGE) (*CCNA_00460-00482*) that is known to spontaneously excise from the *Caulobacter* genome at low frequency (31). This MGE is responsible for biosynthesis of a capsular polysaccharide (31) that is differentially regulated across the cell cycle and confers resistance to the caulophage fCr30 (32). Select genes within this locus have enhanced transcription in Δ*ntrC* (*P* < 10^-5^), including those encoding GDP-L-fucose synthase (*CCNA_00471*), GDP-mannose 4,6 dehydratase (*CCNA_00472*), and a P4 family DNA integrase (*CCNA_00480*) (Table S2). Two NtrC-binding sites also flank a second capsule biosynthesis and regulatory locus (*CCNA_00161-CCNA_00167*) outside of the MGE (Table S3), and deletion of *ntrC* results in significantly enhanced expression of several genes within this locus, including the capsule restriction factor, *hvyA* (32) (3-fold; Table S2). In all cases, NtrC binding sites within the MGE directly overlap reported binding sites of the nucleoid associated protein, GapR (17, 18) and either overlap or are adjacent (within 200 bp) with binding sites for the cell cycle regulators MucR1/2 (19) (Fig 5B and Table S3). Thirty-seven of the 51 total NtrC binding sites that we have identified directly overlap with one of the 599 reported GapR binding sites across the *Caulobacter* genome (18) (Fig 5B and Table S3). *gapR* itself is significantly downregulated by 2- fold in the Δ*ntrC* mutant (Table S2). These results suggest that NtrC has a chromosome structuring role in addition to its direct role in transcriptional regulation of nitrogen assimilation genes.

The promoter region of the cell cycle regulator, *sciP* (*CCNA_00948*) (33, 34), contains an NtrC binding site (Table S3), and the transcription of *sciP* and adjacent flagellar genes, *flgE* and *flgD*, is significantly increased in Δ*ntrC* (Table S2), indicating that NtrC represses transcription from this site. NtrC also directly binds the promoter of *mucR1* (*CCNA_00982*) (Table S3); this regulator, along with SciP, has been implicated in controlling the S®G1 cell cycle transition (19). Like *sciP*, deletion of *ntrC* results in enhanced expression of *mucR1* (Table S2), and we conclude that NtrC also represses transcription from this site. We assessed overlap of NtrC binding sites with SciP binding sites across the genome, but observed no significant overlap (Fig 5B). We note occasional overlap between NtrC and binding sites for the essential cell cycle regulator, CtrA (Fig 5B), including sites within the promoters of *sciP* and *hvyA* (Table S3). An additional cell cycle gene that is regulated by NtrC is *hdaA*, which is reported to inactivate DnaA after replication initiation (35). NtrC binds the chromosome upstream of *hdaA* (Table S3), and deletion of *ntrC* results in significantly diminished transcription of *hdaA* (Table S2). Conversely, the region upstream of the DNA replication inhibitor toxin *socB* (36) (within the *socA* gene) is bound by NtrC (Table S3), and deletion of *ntrC* results in significantly enhanced transcription of *socB* (2-fold) without corresponding induction of the *socA* antitoxin (Table S2). Together, these results provide support for a model in which NtrC can function to modulate expression of key cell cycle/replication regulators in *Caulobacter*.

Transcripts corresponding to the contact-dependent inhibition by glycine zipper proteins (*cdzCDI*) system (37) are highly elevated in Δ*ntrC* relative to WT (15-22-fold) (Fig 4A and Table S2), although the nearest NtrC ChIP-seq peak resides downstream of the promoter of this operon, within *cdzI*, itself (Fig 4D and Table S3). It is unclear whether expression of these genes is directly impacted by NtrC, but this NtrC binding site overlaps with a reported GapR binding site (18). *CCNA_02727,* encoding an un-characterized PhoH family protein (38, 39), provides yet another example of gene with overlapping NtrC and GapR binding sites (18) in its promoter that exhibits strongly increased transcription in Δ*ntrC* relative to WT (10-fold) (Fig 4A&D and Fig 7A-B).

### Glutamine and *glnBA* activation rescue the Δ*ntrC* transcriptional defect

Glutamine supplementation res-cued the growth defect of Δ*ntrC* in PYE complex medium (Fig 2B), which raised the question of whether glutamine supplementation would also restore the global transcriptional defect of Δ*ntrC* in PYE. Indeed, glutamine supplementation broadly restored transcription of genes dysregulated in the Δ*ntrC* mutant to WT levels (Fig 4B, Table S2, and Fig S6). However, genes directly regulated by NtrC that are involved in nitrogen assimilation remained significantly dysregulated when glutamine was added to the medium (Fig 4B, Table S2, and Fig S6). For example, *glnB* and *glnA* transcript levels remained 15- and 8-fold lower in Δ*ntrC* than in WT in the presence of 9.3 mM glutamine, while *glnK* and *CCNA_01399* remained 6- and 4- fold lower, respectively (Fig 4B and Table S2). Transcripts from the *ntrC* locus, which is autorepressed, also remained significantly elevated in Δ*ntrC*, as did genes of the *cdz* locus (Fig 4B and Table S2).

We further analyzed transcription in the suppressor mutant, Δ*ntrC* P*_glnBA_*::IS3, which permitted us to assess the transcriptome in a strain that lacks *ntrC* but that expresses *glnBA* (Fig 3C). Restoration of *glnBA* expression in this background restored transcription to WT levels for a subset of the loci that were dysregulated in Δ*ntrC*, though transcription of many dysregulated genes was only partially rescued or remained unchanged (Fig 4C, Table S2, and Fig S6). Again, NtrC-regulated genes directly involved in nitrogen assimilation (e.g., *glnK-CCNA_01399*) remained significantly dysregulated in this strain (Fig 4C and Table S2). Furthermore, while *gapR* transcription is significantly reduced in Δ*ntrC*, its transcription is significantly increased (3-fold) above WT in Δ*ntrC* P*_glnBA_*::IS3 to a level that is congruent to WT cultivated in the presence of 9.3 mM glutamine (Table S2). This same effect is observed for the iron-dependent Fur regulon (40) (e.g., *CCNA_00027*, *CCNA_00028*) (Table S2). Thus, for a subset of genes, IS3 insertion at P*_glnBA_* results in a tran-scriptional effect that mimics media supplementation with 9.3 mM glutamine.

### Loss of the *ntrB*-*ntrC* system results in stalk elonga-tion

*Caulobacter* has a dimorphic life cycle wherein each cell division produces two morphologically and developmentally distinct cells including 1) a flagellated, motile swarmer cell and 2) a sessile stalked cell. The *Caulobacter* stalk is a thin extension of the cell envelope and its length is known to be impacted by phosphate limitation (41) and sugar-phosphate metabolism imbalances (42). We observed that Δ*ntrC* mutant cells develop elongated stalks when cultivated in PYE complex medium (Fig 6A- B). Δ*ntrB* and *ntrC*^D56A^ mutants displayed an intermediate stalk elongation phenotype, while stalks of *ntrC*^D56E^ mutants did not differ from WT (Fig 6B). We conclude that loss of *ntrC* function results in development of elongated stalks in complex medium.

**Figure 6.**
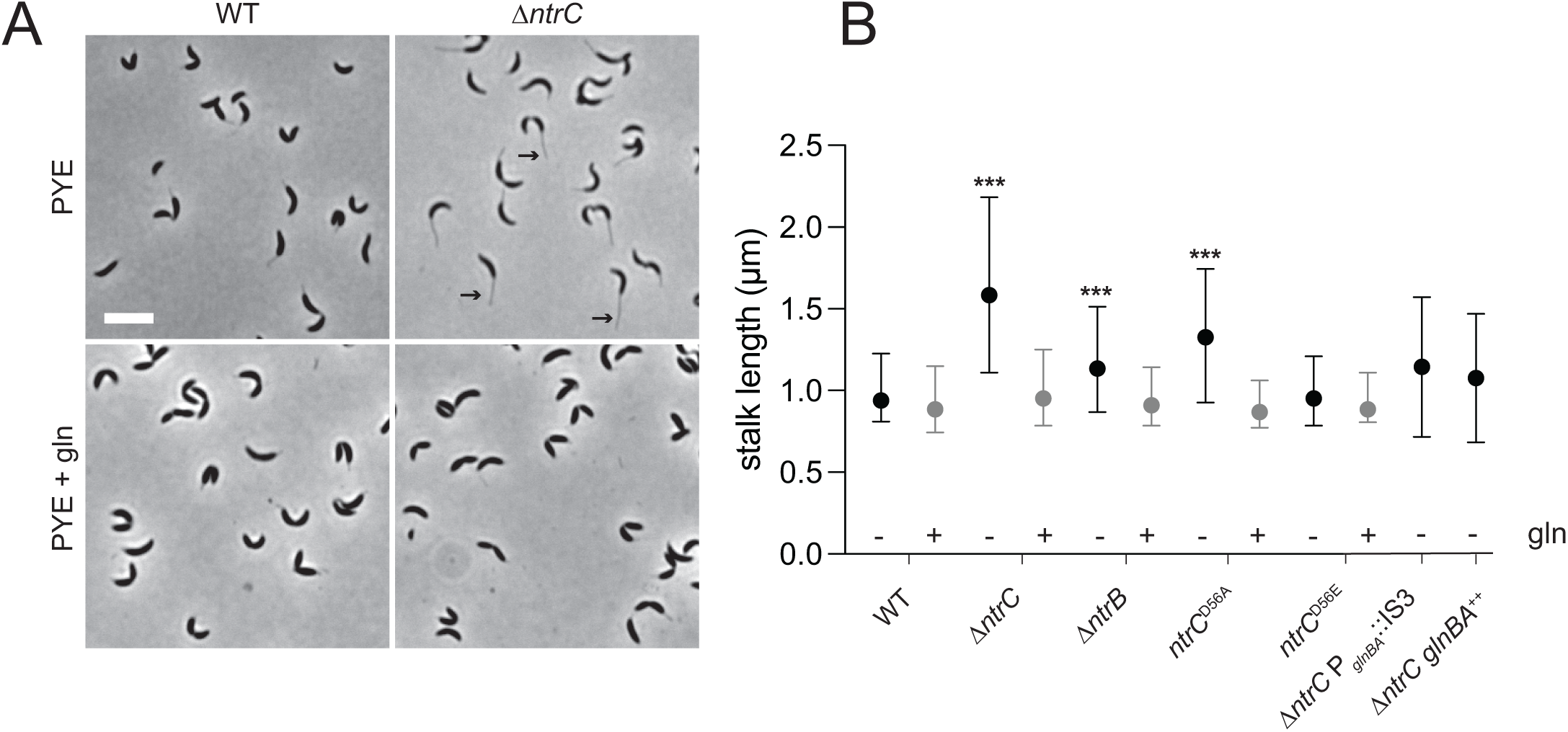
Deletion of the *ntrB-ntrC* two-component system results in development of hyper-elongated stalks. (A) Representative phase-contrast images showing the stalk elongation phenotype of a Δ*ntrC* strain compared to WT; strains were cultivated in PYE complex medium (top). The elongated stalk phenotype is chemically complemented by the addition of 9.3 mM glutamine (gln) to the medium (bottom). Scale bar (white; top left) equals 5 µm. Example stalks in the Δ*ntrC* panel are marked with black arrows. (B) Summary of stalk length measurements for WT, Δ*ntrC*, *ntrC*^D56A^, Δ*ntrB*, *ntrC*^D56A^, Δ*ntrC* P*_glnBA_*::IS3, and Δ*ntrC glnBA*^++^ cultivated without (-/black) and with (+/gray) gln. Data represent median ± interquartile range. Minimum length for stalk segmentation was 0.6 µm. Statistical significance assessed by Kruskal-Wallis test followed by Dunn’s post-test comparison to WT (*** *P* < 0.0001). WT: n=314(- gln) n=207(+ gln); Δ*ntrC*: n=1020(-) n=338(+); Δ*ntrB*: n=440(-) n=75(+); *ntrC*^D56A^: n=849(-) n=204(+); *ntrC*^D56E^: n=339(-) n=177(+); Δ*ntrC* P*_glnBA_*::IS3: n=218(-); Δ*ntrC glnBA*^++^: n=503(-).

Our transcriptomic data showed no evidence of a phosphate limitation response upon *ntrC* deletion, nor did we observe changes in *manA* or *spoT/rsh* expression (Table S2), which have been implicated in stalk elongation (42). We did observe that expression of the *phoH*-like gene, *CCNA_02727*, was elevated 10-fold in Δ*ntrC* compared to WT (Fig 7A). This gene has an NtrC peak in its promoter (Fig 7B), suggesting it is directly repressed by NtrC. PhoH proteins have been implicated in phosphate starvation responses in other bacteria (43, 44), so we tested whether de-repression of this gene in Δ*ntrC* impacted stalk development, which is known to be stimulated by phosphate starvation in *Caulobacter* (41). Overexpression of *CCNA_02727* from a xylose-inducible promoter in WT resulted in significantly longer stalks compared to WT (Fig 7C). However, deletion of *CCNA_02727* in the Δ*ntrC* strain did not ablate stalk elongation (Fig 7C). We conclude that elevated expression of *CCNA_02727* is suffi-cient to promote stalk elongation, but that enhanced ex-pression of *CCNA_02727* in Δ*ntrC* does not solely explain the long stalk phenotype.

**Figure 7.**
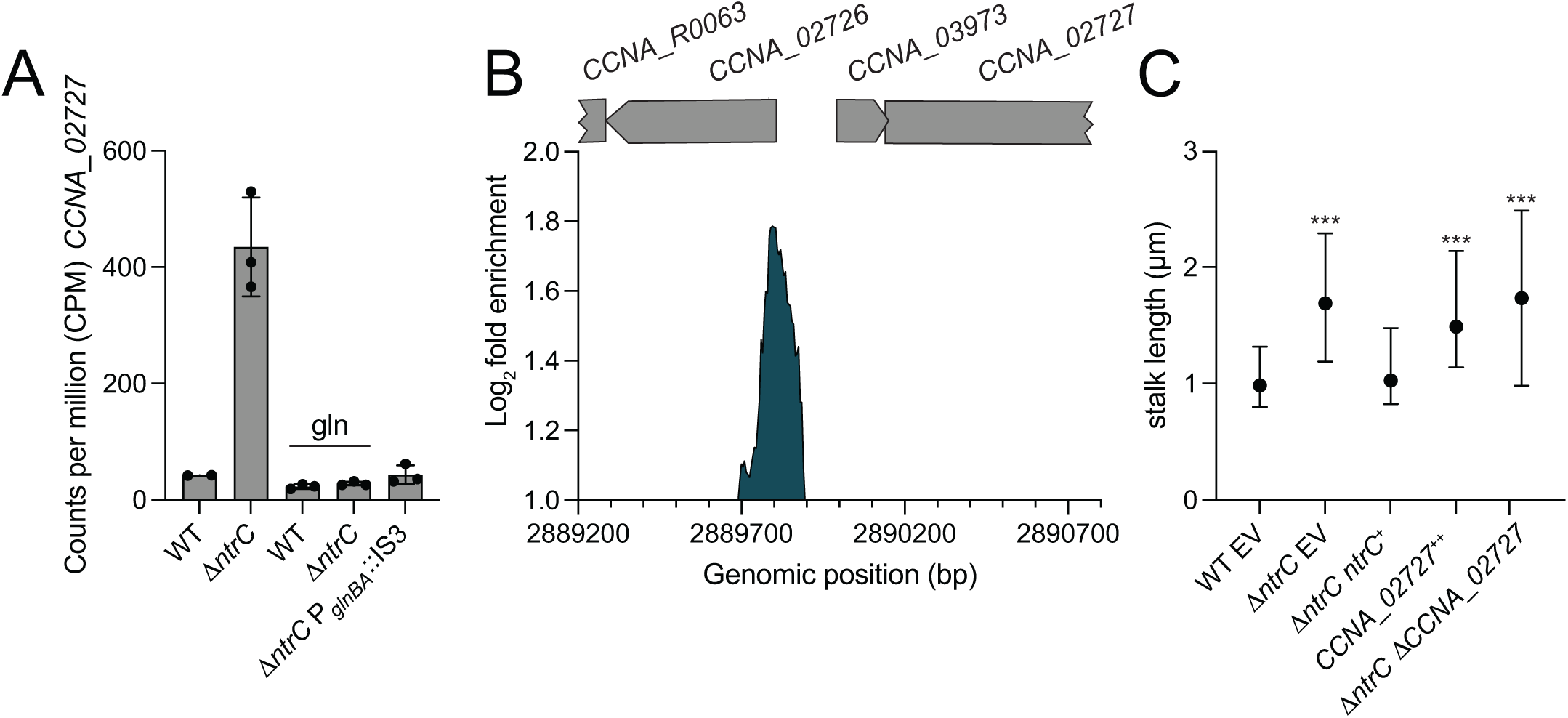
Transcriptional regulation and functional impact of the *phoH*-family gene, *CCNA_02727*. (A) Transcript levels of *CCNA_02727* measured by RNAseq in different genetic backgrounds and conditions: WT and Δ*ntrC* strains grown in PYE complex medium or PYE supplemented with 9.3 mM glutamine (gln), and the Δ*ntrC* P*_glnBA_*::IS3 strain grown in PYE. Data represent mean ± standard deviation of three replicate samples. (B) NtrC chromatin immunoprecipitation sequencing (ChIP-seq) revealed a binding peak upstream of an operon containing the small hypothetical gene, *CCNA_03973*, and *CCNA_02727*. Data represent log_2_ fold enrichment sequence reads in the NtrC immunoprecipitation samples compared to total input sample. Positions of annotated genes are represented by gray bars above the plot. The genomic positions in the reference genome (Genbank accession CP001340) are indicated. (C) Summary of stalk length data, comparing different strains: WT and Δ*ntrC* strains containing an empty vector (EV), a genetic complementation vector (Δ*ntrC ntrC*^+^), Δ*ntrC* Δ*CCNA_02727*, and *CCNA_02727* overexpressed in WT from a xylose-inducible promoter (*CCNA_02727*^++^). The data represent the median and interquartile range; a minimum length of 0.6 µM was used for stalk segmentation. Statistical significance was assessed using the Kruskal-Wallis test followed by Dunn’s test, comparing each condition to WT EV (*** *P* < 0.0001). WT EV: n=330; Δ*ntrC* EV: n=1,481; Δ*ntrC ntrC*^+^: n=366; *CCNA_02727*^++^ n=238; Δ*ntrC* Δ*CCNA_02727*: n=1,261.

Supplementation of PYE with 9.3 mM glutamine fully complemented the stalk length phenotype of Δ*ntrC* (Fig 6A-B) and restoration of *glnBA* expression, either in the suppressor (Δ*ntrC* P*_glnBA_*::IS3) or in the *glnBA* overexpression strain (Δ*ntrC glnBA*^++^), restored Δ*ntrC* stalk length to WT levels (Fig 6B). Similarly, glutamine supplementation complemented stalk length defects of Δ*ntrB* and *ntrC*^D56A^ (Fig 6B). Altogether, these results indicate that stalk elongation phenotype of *ntrB-ntrC* mutants results from an explicit lack of usable nitrogen or a nutrient imbalance due to the reduced availability of usable nitrogen. We note that expression of *CCNA_02727* in Δ*ntrC* is restored to WT levels when PYE is supplemented with glutamine (Fig 7A). This result indicates that regulation of *CCNA_02727* by NtrC is not via a simple, direct repressive mechanism.

### Δ*ntrC* forms mucoid colonies

The *Caulobacter* swarmer and stalked cell types differ not only in cellular morphology, but also in their capsulation state (32). The swarmer cell is non-capsulated, while the stalked cell elaborates an exopolysaccharide (EPS) capsule composed of a repeating tetrasaccharide (45). Capsulation results in enhanced buoyancy which is apparent during centrifugation (32). When centrifuged, Δ*ntrC* cells cultivated in PYE displayed a “soft” or “fluffy” pellet compared to WT (Fig 8A), which suggested that Δ*ntrC* had altered EPS. Over-production of EPS results in colonies that appear mucoid (i.e., glossy) on solid medium containing abundant sugar (31, 32), and Δ*ntrC* displayed a mucoid phenotype on PYE supplemented with 3% sucrose, a condition that has been shown to enhance *Caulobacter* mucoidy (31) (Fig 8B). We conclude that loss of *ntrC* impacts the production or composition of envelope polysaccha-rides.

**Figure 8.**
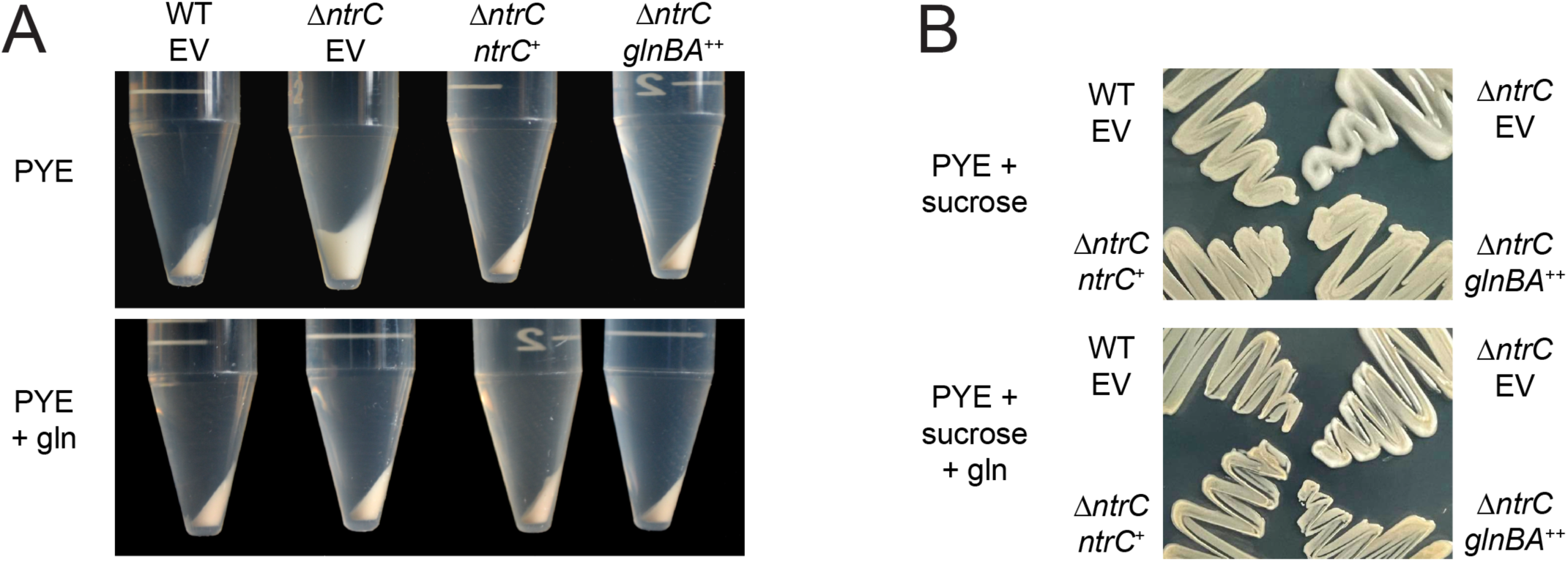
The hyper-mucoid phenotype of Δ*ntrC* in PYE complex medium is suppressed by either glutamine supplementation or *glnBA* expression. (A) Cell pellets of WT and Δ*ntrC* carrying an empty vector (EV) or vectors expressing *ntrC* (*ntrC*^+^) or *glnBA* (*glnBA*^++^). Strains were grown overnight in PYE complex medium or PYE supplemented with 9.3 mM glutamine (gln). Overnight cultures were normalized to OD_660_ = 0.5 and cells from 10 ml were centrifuged at 7,197 x g for 3 min at 4°C, and pellets were photographed. (B) Growth of WT EV, Δ*ntrC* EV, Δ*ntrC ntrC*^+^, and Δ*ntrC glnBA*^++^ on PYE agar supplemented with 3% sucrose (PYE + sucrose) or PYE agar supplemented with 3% sucrose and 9.3 mM glutamine (PYE + sucrose + gln). Plates were incubated for 4 days at 30°C and photographed.

We again tested whether glutamine supplementation could restore a phenotype of Δ*ntrC* to that of WT. Centrifugation of Δ*ntrC* cultures grown in PYE supplemented with 9.3 mM glutamine resulted in a compact pellet like WT (Fig 8A). Furthermore, Δ*ntrC* cultivated on PYE supplemented with sucrose and 9.3 mM glutamine had a WT appearance (Fig 8B). Ectopic expression of *glnBA* in Δ*ntrC* grown in PYE similarly complemented Δ*ntrC* mucoidy (Fig 8A-B). These results support a model in which genetic or chemical restoration of intracellular glutamine levels complements the mucoid phenotype of Δ*ntrC*.

### The Δ*ntrC* mucoidy phenotype requires the MGE

The mucoid appearance of Δ*ntrC* aligns with transcriptomic and ChIP-seq data that show *ntrC*-dependent repression of EPS synthesis genes, including those located within the *Caulobacter* mobile genetic element (MGE) (e.g., *CCNA_00471*, *CCNA_00472*) (Table S2). Considering the numerous NtrC binding sites within the MGE and its role in capsular polysaccharide biosynthesis (31), we tested whether the mucoid phenotype of Δ*ntrC* required the MGE. Specifically, we deleted *ntrC* from a *Caulobacter crescentus* NA1000 strain that had spontaneously lost the MGE (31), resulting in a *Caulobacter* ΔMGE Δ*ntrC* mutant. When cultivated in PYE, the NA1000 ΔMGE Δ*ntrC* strain did not display a “fluffy” pellet or exhibit a mucoid phenotype on solid medium (Fig S7A). We further deleted *ntrC* in *C. crescentus* CB15 strain (CB15 Δ*ntrC*), which similarly lacks the MGE (31). CB15 Δ*ntrC* had WT phenotype in pellet and plate growth assays (Fig S7B). These results provide evidence that the mucoid phenotype of Δ*ntrC* is dependent on the EPS synthesis genes of the MGE. We conclude that transcriptional dysregulation due to loss of NtrC impacts cell envelope polysaccharide production via the MGE and, perhaps, through other genes involved in EPS biosynthesis (Table S3).

## Discussion

### *ntrB-ntrC* differentially impacts growth in defined and complex medium

Environmental nitrogen is an important cell cycle and developmental regulatory cue in *Caulobacter* (8), which motivated us to explore the function of the NtrB-NtrC TCS, a broadly conserved regulator of nitrogen metabolism (6). We characterized the population-level growth phenotypes of *ntrB* and *ntrC* mutants under media conditions containing distinct nitrogen sources and demonstrated that the sensor kinase gene, *ntrB,* and the AAA+-type response regulator gene, *ntrC,* are essential for growth in a defined medium in which NH_4_^+^ is the sole nitrogen source (Fig 1 and Fig 2). Strains expressing a *ntrC* allele harboring a mutated aspartyl phosphorylation site in its receiver domain (*ntrC*^D56A^) also failed to grow in this defined medium (Fig 2G). These data support an expected model in which phosphorylation of NtrC by NtrB is necessary for NH_4_^+^ assimilation. An additional histidine kinase/response regulator pair, *ntrY-ntrX*, is part of the *ntrBC* genetic locus in *Caulobacter* and is postulated to have arisen from gene duplication (25). The inability of Δ*ntrB* to grow in M2G indicates that NtrB (and not NtrY) is the major histidine kinase for NtrC *in vivo.* Each of the three NtrC domains – 1) Receiver, 2) AAA+ ATPase, and 3) HTH DNA binding do-main – are required for growth in ammonium-defined medium (M2G) (Fig 2G). Replacement of NH_4_^+^ with glutamine in M2G fully rescued growth of *ntrC*^ΔHTH^ but not *ntrC*^ΔREC^ and *ntrC*^ΔAAA^ mutants (Fig 2H). This suggests that some component of the growth defect of *ntrC*^ΔREC^ and *ntrC*^ΔAAA^ in defined medium is independent of cellular nitrogen status. Considering the overlap of NtrC binding sites with GapR and MucR1, variants of NtrC without the AAA+ or REC domains might exhibit dominant-negative effects. These truncated alleles could interfere with interactions involving GapR and MucR1, thereby disrupting gene expression at multiple chromosomal loci.

A strain lacking *ntrC* is viable in PYE complex medium but has a reduced growth rate, a phenotype that is complemented by addition of glutamine (Fig 2A-B) (8). Surprisingly, Δ*ntrB* and *ntrC*^D56A^ had no growth rate defect, but did exhibit a growth yield defect in PYE (i.e., final culture density) (Fig 2C) that was rescued by the addition of glutamine to the medium (Fig 2D). From these results, we conclude that NtrC∼P is less important in complex medium during log phase growth and becomes more important at higher cell density when organic nitrogen becomes more limited and waste products accumulate. NtrC domain truncation mutants, *ntrC*^DREC^, *ntrC*^DAAA^, and *ntrC*^DHTH^, grew slower in PYE (Fig 2E), though the *ntrC*^DREC^ and *ntrC*^DAAA^ strains had more severe defects than *ntrC*^DHTH^, which phenocopied Δ*ntrC*. As discussed above, the *ntrC*^DREC^ and *ntrC*^DAAA^ yield phenotypes in PYE may be due to dominant-negative effects of expressing these truncated NtrC polypeptides *in vivo,* though glutamine supplementation to PYE did complement the defects of all NtrC domain truncation mutants in PYE (Fig 2F).

### IS3 transposition repeatedly rescued the growth defect of *ntrC* mutants

The role of NtrC in activating glutamine synthase expression and facilitating ammonium assimilation is well-established in various species (6). We demonstrated that *Caulobacter ntrC* is essential in ammonium-defined medium (M2G) and made the surprising observation that cultures of *Caulobacter* Δ*ntrC* occasionally showed robust growth in M2G; this suggested there was a route for spontaneous genetic rescue of the Δ*ntrC* growth defect. We discovered that these “jackpot”-like cultures (46) were a consequence of random insertion of an IS3-family mobile genetic element at the *glnBA* promoter (P*_glnBA_*) of Δ*ntrC* that restored *glnBA* transcription (Fig 3). IS3 elements are present in multiple copies in the *Caulobacter* NA1000 genome (31), and the IS3-dependent transcriptional rescue phenotype we observe is consistent with a report that IS3 insertion elements can function as mobile promoters (47). We also identified two independent IS3-family (IS511/ISCc3) in-sertions upstream of *glnBA* at nucleotide 2192500 (16 bp upstream of the *glnB* start codon) and nucleotide 2192465 (51 bp upstream of the *glnB* start codon) that rescued growth of *ntrC*^ΔHTH^ mutants in M2G defined medium (Table S1), indicating that this is a facile evolutionary route to rescue loss of *ntrC* function under particular conditions.

*Caulobacter* insertion elements were previously shown to be transcriptionally activated in mutants that accumulate the alarmone (p)ppGpp (48), and Ronneau et al (8) have reported that glutamine limitation results in (p)ppGpp accumulation via activation of the PTS^Ntr^ system in *Caulobacter*. Furthermore, (p)ppGpp accumulates in *Caulobacter* starved for NH_4_^+^ in defined medium (48). We postulate that in the absence of *ntrC,* decreased levels of intracellular glutamine result in (p)ppGpp accumulation and IS3 activation; an NtrC binding peak within an IS3-family element (adjacent to *CCNA_02830*) could contribute to IS3 regulation (Table S3).

### The NtrC regulon in *Caulobacter*: more than just nitrogen metabolism

NtrC binds to multiple sites on the *Caulobacter* chromosome, playing a role in both activating and repressing gene expression. As expected, NtrC directly activates transcription of nitrogen assimilation genes such as *glnBA*, *glnK*, and the putative NH_4_^+^ transporter *CCNA_01399*. Conversely, NtrC represses its own operon demonstrating autoregulation, which is well-established for this class of regulators (49). Our study also identified genes not directly involved in nitrogen assimilation in the NtrC regulon. Nine of the 51 NtrC binding sites are located within a mobile genetic element responsible for biosynthesis of a capsular polysaccharide that is differentially regulated across the cell cycle and confers resistance to a caulophage (32). The impact of *ntrC* on envelope polysaccharide is discussed below.

Thirty-seven of 51 NtrC binding sites (>70%) directly overlap with one of the 599 reported GapR binding sites (18) across the *Caulobacter* genome (Fig 5B and Table S3). GapR is a nucleoid-associated protein that binds positively supercoiled DNA and supports DNA replication (17), suggesting a possible connection between NtrC and chromosome organization/maintenance in *Caulobacter*. In addition, we observed significant overlap in binding sites of NtrC and the cell cycle regulator, MucR1 (19). Beyond *mucR1*, NtrC directly bound upstream and modulated transcription of other genes that impact cell cycle processes, including *sciP*, *hdaA*, and *socB* (33-36). NtrC appears to repress transcription of *sciP* and *mucR1*, which have been implicated in controlling the cell cycle transition from S to G1 upon compartmentalization of the nascent swarmer cell and also represses transcription of *socB*, a DNA replication inhibitor toxin. The exact mechanism of repression at these promoters remains undefined. These findings suggest that NtrC directly impacts regulation of the cell cycle in *Caulobacter*.

NtrC also regulates the *cdzCDI* operon that encodes a bacteriocin cell killing system activated in stationary phase (37). Loss of *ntrC* results in increased expression of the Cdz system; this transcriptional phenotype is not fully complemented by glutamine supplementation to the medium (Fig 4B and Table S2). Thus, repression of this locus by NtrC is not solely determined by nitrogen availability.

### *ntrC* is a stalk elongation factor

*Caulobacter* cell division results in the production of a swarmer cell and a stalked cell. The swarmer cell differentiates into a reproductive stalked cell by shedding its polar flagellum, producing an adhesive holdfast at the same cell pole, and forming and a stalk that extends from that pole. Stalk length is regulated, and phosphate limitation was previously believed to be the only factor that determined *Caulobacter* stalk length (41). However, recent studies have demonstrated that metabolic imbalances in sugar-phosphate metabolism influence stalk elongation (42).

We have demonstrated that stalk elongation is genetically linked to the *ntrB-ntrC* TCS. The deletion of *ntrC*, *ntrB*, or replacement of *ntrC* with a non-phosphorylatable allele (D56A) resulted in hyper-elongated stalks in PYE (Fig 6B). Supplementation of PYE with glutamine or ectopic *glnBA* expression restored stalk lengths of *ntrB* and *ntrC* mutants to WT. We conclude that the stalk lengthening phenotype of *ntrB* and *ntrC* mutants is a consequence of decreased intracellular glutamine and that stalk elongation is linked to loss of *ntrB*-*ntrC* and possibly nitrogen limitation. Notably, limiting NH_4_^+^ in defined growth medium does not result in increased stalk length in *Caulobacter* (50, 51). Furthermore, excess NH_4_^+^ in combination with high pH restricts stalk elongation even when phosphorus is limited (52). These findings indicate that, while a connection between nitrogen availability and stalk length exists, not all nitrogen limitation conditions impact stalk development. Links between nitrogen availability, phosphorus availability, starvation signals such as (p)ppGpp (42), and stalk length are clearly complex and require further research.

Stalk elongation was previously postulated to enhance diffusive surface area, allowing for increased uptake of nutrients (53, 54), but subsequent work indicated that this is unlikely due to diffusion barriers within the stalk (51, 55). A recent model is that stalk lengthening allows *Caulobacter* in surface-attached communities to reach beyond its neighbors to better access available nutrients, thereby outcompeting other attached microbes and assisting in releasing progeny into the environment (52, 55). We predict that when nitrogen becomes limiting in surface-attached communities, the NtrB-NtrC system can cue the cell, perhaps through intracellular glutamine, to lengthen its stalk to better access nitrogen or other nutrients.

NtrC strongly represses transcription of *CCNA_02727*, a gene encoding a PhoH-family protein, and overexpres-sion of *CCNA_02727* in WT cells results in increased stalk length (Fig 7). However, deletion of *CCNA_02727* in a Δ*ntrC* background did not affect the stalk length of Δ*ntrC*. PhoH family proteins typically possess ATPase and ribonuclease activity (38, 39, 44) and are often activated by the Pho regulon under phosphate starvation conditions in bacteria (43, 44). *CCNA_02727* is not regulated by the Pho regulon in *Caulobacter* (56) but is strongly upregulated under other environmental conditions, such as carbon limitation (57) and heavy metal stress (57) in addition to glutamine deprivation via loss of *ntrC* as described here (Fig 7A). Crosstalk between different sensing systems to balance nutrient levels is well described in bacteria (58) and, therefore, it is possible that regulation of *CCNA_02727* has a general role in controlling nutrient balance or stress response in *Caulobacter*.

### *ntrC* regulates envelope polysaccharide production

*Caulobacter* Δ*ntrC* displays a hyper-capsulation phenotype (Fig 8). NtrC orthologs are reported to regulate biofilm formation and EPS production in other bacteria, including *P. aeruginosa*, *V. vulnificus*, and *B. cenocepacia*, where loss of the *ntrB-ntrC* TCS decreases biofilm and EPS production (59-61). In *V. cholerae*, loss of *ntrC* increases biofilm formation and increases expression of EPS gene regulators (21).

Transcriptomic and ChIP-seq data presented in this study identified an NtrC peak in the promoter of *hvyA*, a gene encoding a transglutaminase homolog that prevents capsulation of swarmer cells (32). Although deletion of *hvyA* increases *Caulobacter* capsulation, its transcription is increased in Δ*ntrC* by 3-fold. The link between *hvyA* expression and the Δ*ntrC* capsule/mucoid phenotype, if any, remains undefined. We further observed an NtrC peak in the promoter region of the operon containing the *CCNA_00471* (*fcI*; GDP-L-fucose synthase) and *CCNA_00472* (GDP-mannose 4,6 dehydratase) genes (Table S3), which reside in the MGE of the *Caulobacter* NA1000 genome. The transcription of these two genes increased 2-fold and 3-fold in Δ*ntrC* relative to WT, respectively. These enzymes function in the two-step synthesis of fucose, which is one of the sugars comprising the tetra-saccharide capsule of *Caulobacter*. It is reported that loss of these genes leads to a significant reduction in EPS production (62). The upregulation of *CCNA_00471-00472* in Δ*ntrC* may contribute to an increase in EPS production and, consequently, the hyper-mucoid and buoyancy phenotypes of Δ*ntrC*. This is supported by the observation that *Caulobacter* Δ*ntrC* strains lacking the MGE (i.e., NA1000 ΔMGE Δ*ntrC*; CB15 Δ*ntrC*) are not mucoid (Fig S7). However, EPS production is a complex process that involves multiple pathways and other genetic and physiological factors could also contribute to the envelope polysaccharide phenotype of Δ*ntrC*. Indeed, EPS production is apparently linked to changes in intracellular glutamine levels independent of NtrC, given that adding glutamine to the medium represses EPS gene expression in Δ*ntrC* (Table S2). The effect of glutamine on EPS gene transcription is congruent with our observation that either addition of glutamine to PYE or the ectopic expression of *glnBA* complements the mucoid phenotype of Δ*ntrC* (Fig 8).

### An unconventional NtrC

*Caulobacter* NtrC is lacks a GAFTGA motif within its primary structure (Fig S1), which is necessary for interaction with s^54^ (13). Consistent with previous reports of NtrC orthologs lacking a GAFTGA motif (14, 15), our data indicate that NtrC regulates s^70^-dependent promoters. For example, NtrC-repressed genes such as *hvyA* and *sciP* are activated by CtrA, a s^70^-dependent transcriptional regulator (32-34). The NtrC binding peak summits within P*_hvyA_* and P*_sciP_* reside 4 bp and 55 bp from the CtrA peak summit at these promoters, respectively (Table S3), indicating that NtrC may directly compete with CtrA at these sites to repress transcription. We also identified NtrC-activated genes that possess s^70^ promoters such as *hdaA*, which is also activated by DnaA (35), a s^70^-dependent regulator (63, 64). The mechanism by which *Caulobacter* NtrC functions at s^70^ promoters remains unclear.

Mutation of the conserved NtrC aspartyl phosphorylation site (D56) results in reduced transcriptional activation of the *glnBA* locus (Fig S4), highlighting the important role of this residue in NtrC-mediated transcriptional activation (at *glnBA*). Similarly, in *R. capsulatus* NtrC, which also lacks GAFTGA, aspartyl phosphorylation is required for transcriptional activation (14, 15). *V. cholerae* VspR, a bEBP that lacks GAFTGA and regulates s^70^ promoters, does not require phosphorylation but utilizes the conserved aspartyl phosphorylation site for phosphate sensing (16). Whether D56 phosphorylation differentially affects NtrC function at binding sites across the *Caulobacter* chromosome is not known. In *R. capsulatus* NtrC, ATP binding rather than hydrolysis by the AAA+ domain is essential for transcriptional activity (15), while VspR does not require ATP to function (65). We have shown that conserved residues of the Walker A and Walker B motifs in the *Caulobacter* NtrC AAA+ domain are required for NH_4_^+^ utilization in defined medium (Fig S3), providing evidence that ATP binding and ATP hydrolysis by NtrC are necessary for controlling the gene expression program that underlies NH_4_^+^ assimilation. More generally, we predict that ATP binding and hydrolysis by *Caulobacter* NtrC contribute to the regulation of σ^70^-dependent promoters, distinguishing it from other unconventional σ^70^-regulating bEBPs. The *Caulobacter* genome (31) encodes four bEBPs: NtrC, NtrX, FlbD, and TacA. Unlike NtrC and NtrX, FlbD and TacA possess the GAFTGA motif. Notably, TacA regulates stalk biogenesis by controlling expression of s^54^-dependent genes, including *staR* (66). Our study establishes a genetic link between the *ntrB-ntrC* TCS and the *Caulobacter* stalk. Thus, development of the polar stalk structure is controlled by at least two distinct bEBPs, NtrC and TacA, which are regulated by different environmental stimuli and have distinct primary structural and regulatory properties.

## Materials and Methods

### Growth conditions

*E. coli* strains were cultivated in Lysogeny Broth (LB) [10 g tryptone, 5 g yeast extract, 10 g NaCl per L] or LB solidified with 1.5% (w/v) agar at 37˚C. LB was supplemented with appropriate antibiotics when necessary. Antibiotic concentrations for selection of *E. coli* were as follows: kanamycin 50 µg/ml, chloramphenicol 20 µg/ml, carbenicillin 100 µg/ml. *Caulobacter* strains were cultivated in peptone yeast extract (PYE) [2 g/L peptone, 1 g/L yeast extract, 1 mM MgSO_4_, 0.5 mM CaCl_2_] complex medium or PYE solidified with 1.5% (w/v) agar at 30˚C or 37˚C. Antibiotic concentrations for selection of *Caulobacter* were as follows: kanamycin 25 µg/ml (in solid medium), 5 µg/ml (in liquid medium), chloramphenicol 1.5 µg/ml. Nalidixic acid (20 µg/ml) was added to counterselect *E. coli* after conjugations. For glutamine supplementation experiments in PYE, 9.3 mM (final concentration) glutamine was added. For experiments in defined medium, *Caulobacter* strains were grown in M2 minimal salts medium with glucose (M2G) [6.1 mM Na_2_HPO_4_, 3.9 mM KH_2_PO_4_, 9.3 mM NH_4_Cl, 0.25 mM CaCl_2_, 0.5 mM MgSO_4_, 10 uM ferrous sulfate chelated with EDTA (Sigma), and 0.15% glucose]. For glutamine supplementation experiments in M2G, NH_4_^+^ was replaced with molar-equivalent (9.3 mM final concentration) glutamine.

### Strains and plasmids

Strains, plasmids, and primers used in this study are presented in Table S4. To generate plasmid constructs for in-frame deletions and other allele replacements, homologous upstream and downstream fragments (∼500 bp/each) were PCR-amplified and joined via overlap extension PCR (67). PCR products were cloned into plasmid pNPTS138 by restriction enzyme digestion and ligation. Similarly, to create genetic complementation constructs, target genes were amplified and fused to their upstream promoters (∼500 bp fragment immediately upstream of the start of the annotated operon) via overlap extension PCR and these fused PCR products were purified and cloned into pXGFPC-2 (pMT585) (68), a plasmid that in-tegrates into the *xylX* locus in *Caulobacter*. For complementation, the genes with their native promoters were cloned in the opposite orientation of the P*_xylX_* promoter in this plasmid. For xylose-inducible expression, target genes were PCR-amplified and ligated into pMT585 in the same orientation as (i.e., downstream of) the P*_xylX_* pro-moter. To create the *glnBA* transcriptional reporter construct, the target promoter (∼500 bp fragment upstream of the start of the *glnBA* operon) was PCR-amplified and cloned into pPTM056 (69), which resulted in the fusion of P*_glnBA_* to *mNeonGreen*. All ligations were transformed into *E. coli* TOP10. All plasmids were sequence confirmed.

Plasmids were transformed into *Caulobacter* via electroporation or triparental mating from TOP10 using FC3 as a helper strain (70). In-frame deletion and allele replacement strains were generated via two-step recombination using *sacB* counterselection using an approach similar to that described by Hmelo et al (71). Briefly, primary recombinants bearing pNPTS138-derived allele-replacement plasmids were selected on solidified PYE containing kanamycin. Single colonies were then grown in PYE broth without selection for 6-18 hours (h) before secondary recombinants were selected on PYE containing 3% sucrose. The resulting clones were screened to confirm kanamycin sensitivity. Then allele replacement was confirmed by PCR for in-frame deletion alleles or PCR amplification and Sanger sequencing for point mutation alleles.

### Measurement of growth in PYE complex medium

Starter cultures were grown overnight in PYE or PYE plus 9.3 mM glutamine shaken at 30˚C. Overnight cultures were diluted to OD_660_ 0.1 in the same media and incubated shaking for 2 h at 30˚C to bring cultures to a similar (logarithmic) phase of growth. Cultures were then diluted to OD_660_ 0.025 in the same media and shaken at 30˚C. Optical density at 660 nm was measured at the timepoints indicated.

### Measurement of growth in M2G defined medium

Starter cultures were shaken overnight in PYE at 30˚C. Starter cultures were pelleted and washed three times with M2G or M2G in which NH_4_^+^ was replaced with molar equivalent glutamine (9.3 mM final concentration) before dilution to OD_660_ 0.025 in the respective medium. These cultures were incubated at 30 ˚C with shaking for 24 h and culture density was measured optically (OD_660_).

### Selection of Δ*ntrC* and *ntrC*^ΔHTH^ suppressors

When Δ*ntrC*, *ntrC*^ΔHTH^, or *ntrC*^D56A^ cultures incubated in M2G overnight at 30˚C exhibited visible turbidity, cultures were spread on PYE to isolate individual colonies bearing suppressing mutations. These putative suppressor strains were re-inoculated into M2G to confirm growth in the absence of a functional *ntrC* allele. Strains that grew rapidly - similar to WT - were saved and genomic DNA was purified and sequenced. Briefly, genomic DNA was extracted from 1 ml of saturated PYE culture using guanidinium thiocyanate (72). Genomic DNA was sequenced (150 bp paired-end reads) at SeqCenter (Pittsburgh, PA) using an Illumina NextSeq 2000. DNA sequencing reads were mapped to the *Caulobacter* NA1000 genome (Genbank accession CP001340) (31) and polymorphisms were identified using breseq (73).

### RNA extraction, sequencing, and analysis

Starter cultures were grown for 18 h at 30˚C in PYE or PYE plus 9.3 mM (final concentration) glutamine. Cultures were then diluted to OD_660_ 0.1 in their respective medium and grown for 2 h to get the cultures in similar (logarithmic) phase of growth. Once again, cultures were diluted to OD_660_ 0.1 in their respective medium and grown another 3.25 h (OD_660_ < 0.4) to capture mRNA in a similar log phase of the growth curve. 6 ml of each culture were pelleted via centrifugation (1 min at 17,000 x g). Pellets were immediately resuspended in 1ml TRIzol and stored at -80˚C until RNA extraction. To extract RNA, thawed samples were incubated at 65˚C for 10 min. After addition of 200 µl of chloroform, samples were vortexed for 20 s and incubated at room temperature (RT) for 5 min. Phases were separated by centrifugation (10 min at 17,000 x g). The aqueous phase was transferred to a fresh tube and an equal volume of isopropanol was added to precipitate the nucleic acid. Samples were stored at 80˚C (1 h to overnight), then thawed and centrifuged at 17,000 x g for 30 min at 4˚C to pellet the nucleic acid. Pellets were washed with ice-cold 70% ethanol then centrifuged for at 17,000 x g for 5 min at 4˚C. After discarding the supernatant, pellets were air-dried at RT, resuspended in 100 µl RNAse-free water, and incubated at 60˚C for 10 min. Samples were treated with TURBO DNAse (Invitrogen) following manufactures protocol for 30 min at RT and then column purified using RNeasy Mini Kit (Qiagen). RNA samples were sequenced at SeqCenter (Pittsburgh, PA). Briefly, sequencing libraries were prepared using Illumina’s Stranded Total RNA Prep Ligation with Ribo-Zero Plus kit and custom rRNA depletion probes. 50 bp paired end reads were generated using the Illumina NextSeq 2000 platform (Illumina). RNA sequencing reads are available in the NCBI GEO database under series accession GSE234097. RNA sequencing reads were mapped to the *Caulobacter* NA1000 genome (31) using default mapping parameters in CLC Genomics Workbench 20 (Qiagen). To identify genes regulated by NtrC, the following criteria were use: fold change > 1.5, FDR *P* < 0.000001 and maximum group mean RPKM > 10. Gene expression data were hierarchically clustered in Cluster 3.0 (74) using an uncentered correlation metric with average linkage. The gene expression heatmap was generated using Java TreeView (75).

### Chromatin immunoprecipitation with sequencing (ChIP-seq)

*Caulobacter ntrC* was PCR-amplified and inserted into pPTM057-3xFLAG expression vector via restriction digestion and ligation to generate a 3xFLAG-NtrC fusion expressed from a cumate-inducible promoter. This suicide plasmid was propagated in *E. coli* TOP10 and conjugated into *Caulobacter* Δ*ntrC* to integrate at the xylose locus. For ChIP-seq experiments, the Δ*ntrC xylX*::pPTM057- 3xFLAG-*ntrC* strain was grown overnight in PYE at 30˚C. The overnight culture was diluted to OD_660_ 0.1 in PYE and outgrown for 2 h at 30˚C. This culture was back-diluted to OD_660_ 0.1 in PYE supplemented with 50 µM cumate and grown for 3.25 h at 37˚C to induce 3xFLAG-*ntrC* during log phase growth. To crosslink 3xFLAG-NtrC to DNA, formaldehyde was added to 125 ml of culture to a final concentration of 1% (w/v) and shaken at 37˚C for 10 min. The crosslinking was quenched using a final concentration of 125 mM glycine and shaken at 37˚C for 5 min. Cells were pelleted by centrifugation at 7,196 x g for 5 min at 4˚C. Supernatant was removed and the pellet was washed 4 times with ice-cold PBS pH 7.5. To lyse the cells, the washed pellet was resuspended in 1 ml lysis buffer [10 mM Tris pH 8, 1 mM EDTA, protease inhibitor tablet (Roche), 1 mg/ml lysozyme]. After a 30 min incubation at 37˚C for 0.1% (w/v) sodium dodecyl sulfate (SDS) was added. To shear the genomic DNA to 300-500 bp fragments, the lysate was sonicated on ice for 10 cycles (20% magnitude for 20 sec on/off pulses using a Branson Sonicator). Cell debris was cleared by centrifugation (15,000 x g for 10 min at 4°C). Supernatant was transferred to a clean tube and Triton X-100 was added to a final concentration of 1% (v/v). The sample was pre-cleared via incubation with 30 µl of SureBeads Protein A magnetic agarose beads (BioRad) for 30 min at RT. The supernatant was transferred to a clean tube and 5% of the total lysate was saved as the input DNA reference sample. Pulldown was performed as previously described (69). Briefly, 100 ul magnetic agarose anti-FLAG beads (Pierce / Thermo) were pre-equilibrated in binding buffer [10 mM Tris pH 8 at 4°C, 1 mM EDTA, 0.1% (w/v) SDS, 1% (v/v) Triton X-100] supplemented with 1% (w/v) bovine serum albumin (BSA) overnight at 4˚C, washed with binding buffer and incubated in the lysate for 3 h at RT. Beads were cleared from the lysate with a magnet, and washed with a low-salt buffer [50 mM HEPES pH 7.5, 1% (v/v) Triton X-100, 150 mM NaCl], followed by a high-salt buffer [50 mM HEPES pH 7.5, 1% (v/v) Triton X-100, 500 mM NaCl], and then LiCl buffer [10 mM Tris pH 8 at 4°C, 1 mM EDTA, 1% (w/v) Triton X-100, 0.5% (v/v) IGEPAL CA-630, 150 mM LiCl]. Finally, beads were incubated with 100 ul elution buffer [10 mM Tris pH 8 at 4°C, 1 mM EDTA, 1% (w/v) SDS, 100 ng/μl 3xFLAG peptide] for 30 min at RT. After pulldown, the input sample was brought to equal volume as the output/pulldown sample using elution buffer [10 mM Tris pH 8, 1 mM EDTA pH 8, 1% SDS, 100 ng/µl 3xFLAG peptide]. Input and output samples were supplemented with 300 mM NaCl and 100 µg/ml RNAse A and incubated at 37°C for 30 min. Proteinase K was added to samples at a final concentration of 200 µg/ml and samples were incubated overnight at 65°C to reverse crosslinks. Samples were purified using the Zymo ChIP DNA Clean & Concentrator kit. ChIP DNA was sequenced at SeqCenter (Pittsburgh, PA). Briefly, sequencing libraries were prepared using the Illumina DNA prep kit and sequenced (150 bp paired end reads) on an Illumina Nextseq 2000. ChIP-seq sequence data have been deposited in the NCBI GEO database under series accession GSE234097.

### ChIP-seq analysis

Paired-end reads were mapped to the *C*. *crescentus* NA1000 reference genome (GenBank accession number CP001340) with CLC Genomics Workbench 20 (Qiagen). Peak calling was performed with the Genrich tool (https://github.com/jsh58/Genrich) on Galaxy; peaks are presented in Table S3. Briefly, PCR duplicates were removed from mapped reads, replicates were pooled, input reads were used as the control dataset, and peak were called using the default peak calling option [Maximum q-value: 0.05, Minimum area under the curve (AUC): 20, Minimum peak length: 0, Maximum distance between significant sites: 100].

To identify promoters that contained NtrC peaks, promoters were designated as 300 bp upstream and 100 bp downstream of the transcription start sites (TSS) annotated for each operon (76, 77). For genes/operons that did not have an annotated TSS, the +1 nucleotide of the first gene in the operon was designated as the TSS. Promoters were defined as containing an NtrC peak if there was any overlap between the NtrC ChIP-seq peak and the indicated promoter. To compare the relative location of NtrC binding sites with various cell cycle regulators, ChIP-peakAnno (78) was used to determine distance from the summit of the NtrC peaks to the nearest CtrA, SciP, MucR1, and GapR peak summit. To compare the relative location of NtrC binding sites with various cell cycle regulators, ChIPpeakAnno (78) was used to determine distance from the summit of the NtrC peaks to the nearest CtrA, SciP, MucR1, and GapR peak summit. ChIP-seq peaks (50 bp windows) for CtrA, SciP, and MucR1 were derived from (19) and the summits were considered the center of the 50 bp window. ChIP-seq summits for GapR were derived from (18). For motif discovery, sequences of the ChIP-seq peaks were submitted to MEME suite (79). Sequences were scanned for enriched motifs between 6 and 30 bp in length that had any number of occurrences per sequence.

### NtrC protein purification

*Caulobacter ntrC* was PCR-amplified and inserted into a pET23b-His6-SUMO expression vector using classical restriction digestion and ligation, such that ntrC was inserted 3’ of the T7 promoter and the His6-SUMO coding sequence. After sequence confirmation, pET23b-His6-SUMO-*ntrC* was transformed into chemically competent *E. coli* BL21 Rosetta (DE3) / pLysS. This strain was grown in 1 L of LB at 37˚C. When the culture density reached approximately OD_600_ ≈ 0.4, expression was induced with 0.5 mM isopropyl β-D-1-thiogalactopyranoside (IPTG) overnight at 16˚C. Cells were harvested by centrifugation (10,000 x g for 10 min) and resuspended in 20 ml lysis buffer [20 mM Tris pH 8, 125 mM NaCl, 10 mM imidazole] and stored at -80˚C until purification.

For protein purification, resuspended cell pellets were thawed at RT. 1 mM phenylmethylsulfonyl fluoride (PMSF) was added to inhibit protease activity and DNase I (5 µg/ml) was added to degrade DNA after cell lysis. Cells incubated on ice were lysed by sonication (Branson Instruments) at 20% magnitude for 20 sec on/off pulses until the suspension was clear. The lysate was cleared of cell debris by centrifugation (30,000 x g for 20 min) at 4˚C. The cleared lysate was applied to an affinity chromatography column containing Ni-nitrilotriacetic acid (NTA) superflow resin (Qiagen) pre-equilibrated in lysis buffer. Beads were washed with wash buffer [20 mM Tris pH 8, 125 mM NaCl, 30 mM imidazole]. Protein was eluted with elution buffer [20 mM Tris pH 8, 125 mM NaCl, 300 mM imidazole]. The elution fractions containing His6-SUMO-NtrC (∼52 kDa) were pooled and dialyzed in 2 L dialysis buffer [20 mM Tris pH8, 150 mM NaCl] for 4 h at 4˚C to dilute the imidazole. Purified ubiquitin-like-specific protease 1 (Ulp1) was added to the eluted His6-SUMO-NtrC containing solution which was then dialyzed overnight at 4˚C in 2 L fresh dialysis buffer to cleave the His6-SUMO tag. Digested protein was mixed with 3 ml of NTA superflow resin (Qiagen) that had been pre-equilibrated in wash buffer. After incubation for 30 min at 4˚C, the solution was placed onto a gravity drip column at 4˚C. Flowthrough containing cleaved NtrC was collected and used to generate α-NtrC polyclonal antibodies (Pacific Immunology).

### Western blotting

To prepare cells for analysis, overnight PYE cultures of *Caulobacter* strains in Fig S2B and S2C were diluted in fresh PYE to OD_660_ 0.1 and grown 2 h at 30˚C. These outgrown cultures were then re-diluted in fresh PYE to OD_660_ 0.1 and grown for 3.25 h at 30˚C to capture exponential growth phase. Cells from 1 ml of each culture were collected by centrifugation (12,000 x g for 1 min). After discarding the supernatant, cell pellets were stored at - 20˚C until western blot analysis. Strains in Fig S2A were grown as above except that the outgrowth medium was supplemented with 0.15% xylose and upon re-dilution in xylose supplemented medium, cultures were grown for 24 h at 30˚C to capture stationary growth phase (OD_660_ > 0.6). Cells from 1 ml of each stationary phase culture were harvested as above and stored at -20˚C until western blot analysis. Strains for Fig 2SD were grown in PYE overnight. Cells from 1 ml of each overnight culture were collected by centrifugation as described above and the pellets were placed at -20˚C until western blot analysis.

For western blot analysis, cell pellets were thawed and resuspended in 2X SDS loading buffer [100 mM Tris-Cl (pH 6.8), 200 mM dithiothreitol, 4% (w/v) SDS, 0.2% bromophenol blue, 20% (v/v) glycerol] to a concentration of 0.0072 OD_660_ • ml culture / µl loading buffer. After resuspension, genomic DNA is digested by incubation with 1 µl Benzonase per 50 µl sample volume for 20 min at RT. Samples then were denatured at 95˚C for 5 min. 10 µl of each sample was loaded onto a 4-20% mini-PROTEAN precast gel (Bio-Rad) (Fig S2C) or a 7.5% mini-PROTEAN precast gel (BioRad) (Fig S2A, S2B, and S2D) and resolved at 180 V at RT. Separated proteins were transferred from the acrylamide gel to a PVDF membrane (Millipore) using a semi-dry transfer apparatus (BioRad) at 10 V for 30 min at RT [1X Tris-Glycine, 20% methanol]. The membrane was blocked in 10 ml Blotto [1X Tris-Glycine with 0.1% Tween 20 (TBST) + 5% (w/v) powdered milk] for 1 h to overnight at 4˚C. The membrane was then incubated in 10 ml Blotto + polyclonal rabbit α-NtrC antiserum (1:1,000 dilution) 1 h to overnight at 4˚C. The membrane was washed in TBST three times. The membrane was then incubated in 10 ml Blotto + goat α-rabbit poly-horse-radish peroxidase secondary antibody (Invitrogen; 1:10,000 dilution) for 1-2 h at RT. The membrane was then washed three times with TBST and developed with ProSignal Pico ECL Spray (Prometheus Protein Biology Products). Immediately upon spraying, the membrane was imaged using BioRad ChemiDoc Imaging System (BioRad).

### *Caulobacter* stalk length measurement and analysis

To prepare stationary phase cells, starter cultures were grown in PYE overnight at 30˚C and diluted to OD_660_ 0.1 in fresh PYE or PYE plus 9.3 mM glutamine. After a 2 h outgrowth at 30˚C cultures were re-diluted to OD_660_ 0.1 in fresh medium and grown for 24 h at 30˚C to capture stalk lengths in stationary phase (> OD_660_ 0.6). 2 µl of each stationary phase culture were spotted on an agarose pad [1% agarose dissolved in water] on a glass slide and covered with a glass cover slip. Cells were imaged using a Leica DMI 6000 microscope using phase contrast with an HC PL APO 63x/1.4 numeric aperture oil Ph3 CS2 objective. Images were captured with an Orca-ER digital camera (Hamamatsu) controlled by Leica Application Suite X (Leica). Stalk length was measured using BacStalk (80) with a minimum stalk length threshold of 0.6 microns.

### Transcriptional reporter assay

Overnight starter cultures grown in PYE supplemented with chloramphenicol (1.5 µg/ml) to maintain the replicating plasmid were diluted to OD_660_ 0.1 in the same medium and outgrown for 2 h at 30˚C. Outgrown cultures were re-diluted to OD_660_ 0.1 in the same medium and grown at 30˚C for 24 h to capture expression in stationary phase. 200 µl of each culture was transferred to a Costar flat bottom, black, clear bottom 96-well plate (Corning). Cell density assessed by absorbance (660 nm) and mNeonGreen fluorescence (excitation = 497 ± 10 nm; emission = 523 ± 10 nm) were measured in a Tecan Spark 20M plate reader.

## Acknowledgements

We thank members of the Crosson Lab for helpful feed-back over the course of this study. Research reported in this publication was supported in part by the National Institute of General Medical Science of the National Institutes of Health under award number R35GM131762 and by Army Research Office contract W911NF2210105 to S.C.

## Supplemental Figures

**Figure S1.**
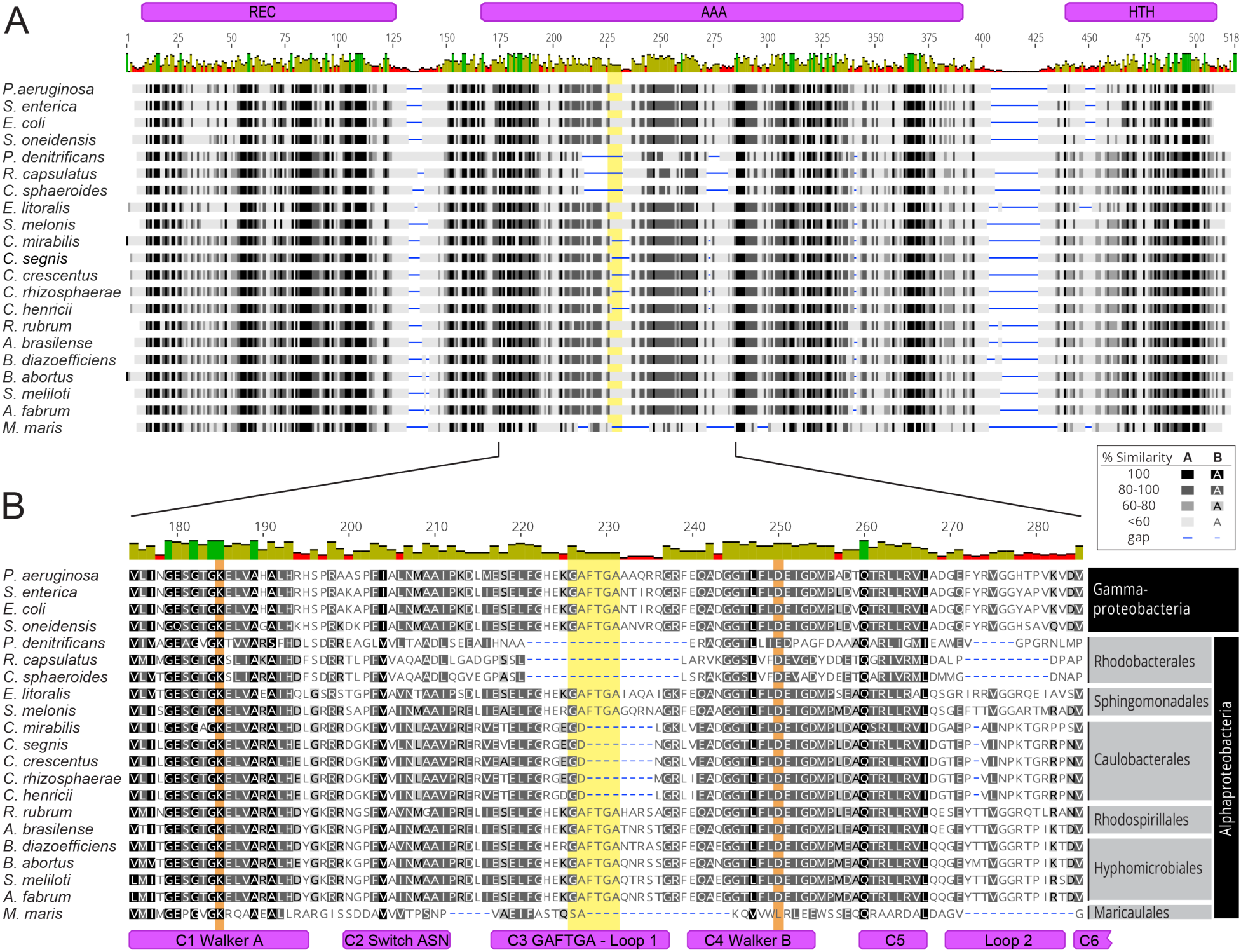
Multiple sequence alignment revealing conserved and distinctive characteristics of NtrC/GlnG se-quences. (A) Overview of full length NtrC/GlnG protein sequence alignment. Residue numbers indicate alignment positions. (B) Highlight of the sequences within the AAA+ domain including the Walker A, Walker B and GAFTGA loop regions. Alignment was performed using Clustal Omega {McWilliam, 2013 #77}, and similarity scores were calculated based on the Blosum62 matrix. In the alignment, gaps are denoted by blue lines. The L1 loop containing the GAFTGA motif is highlighted in yellow. Key ATP binding and ATP hydrolysis residues described in the main text are highlighted in orange. The domains are described following the nomenclature outlined in Bush and Dixon {Bush, 2012 #16}. Spe-cies are indicated at the left of aligned sequences. Full organism names and genbank accessions for this analysis in the order presented in the alignment are: *Pseudomonas aeruginosa* PAO1 (NZ_CP053028), *Salmonella enterica* subsp. enterica serovar Typhimurium str. 798 (NC_017046), *Escherichia coli* str. K-12 substr. MG1655 (NZ_CP027060), *She-wanella oneidensis* MR-1 (NC_004347), *Paracoccus denitrificans* PD1222 (NC_008687), *Rhodobacter capsulatus* SB 1003 (NC_014034), *Cereibacter sphaeroides* 2.4.1 (formerly *Rhodobacter sphaeroides*; NC_007493), *Erythrobacter litoralis* DSM 8509 (NZ_CP017057), *Sphingomonas melonis* TY (NZ_CP017578), *Caulobacter mirabilis* (NZ_CP024201), *Caulobacter segnis* ATCC 21756 (NC_014100), *Caulobacter crescentus* NA1000 (NC_011916), *Cau-lobacter rhizosphaerae* (NZ_CP048815), *Caulobacter henricii* (NZ_CP013002), *Rhodospirillum rubrum* ATCC 11170 (NC_007643), *Azospirillum brasilense* (NZ_CP012915), *Bradyrhizobium diazoefficiens* USDA 110 (formerly B. japoni-cum; NC_004463), *Brucella abortus* 2308 (NC_007618), *Sinorhizobium meliloti* 1021 (NC_003037), *Agrobacterium fabrum* str. C58 (formerly A. tumefaciens; NC_003062), *Maricaulis maris* MCS10 (NC_008347). We note that within the Caulobacterales, we also searched the following complete genomes: *Asticcacaulis excentricus* CB 48 (NC_014816), *Brevundimonas diminuta* (NZ_CP021995), *Brevundimonas nasdae* (NZ_CP080036) and *Brevundimo-nas subvibrioides* ATCC 15264 (NC_014375). We found that genomes from the genera *Asticcacaulis* and *Brevundimo-nas* lack *ntrC* and possess only *ntrX* at the *ntr* locus. NtrX appears to be specific to the Alphaproteobacteria and lacks the classical GAFTGA sequence motif; NtrX sequences exhibit a shorter motif (EG**EG---G**xxK/R) where the dashes are gaps compared to the motif found in most enhancer binding proteins (EK**GAFTGA**xxK/R).

**Figure S2.**
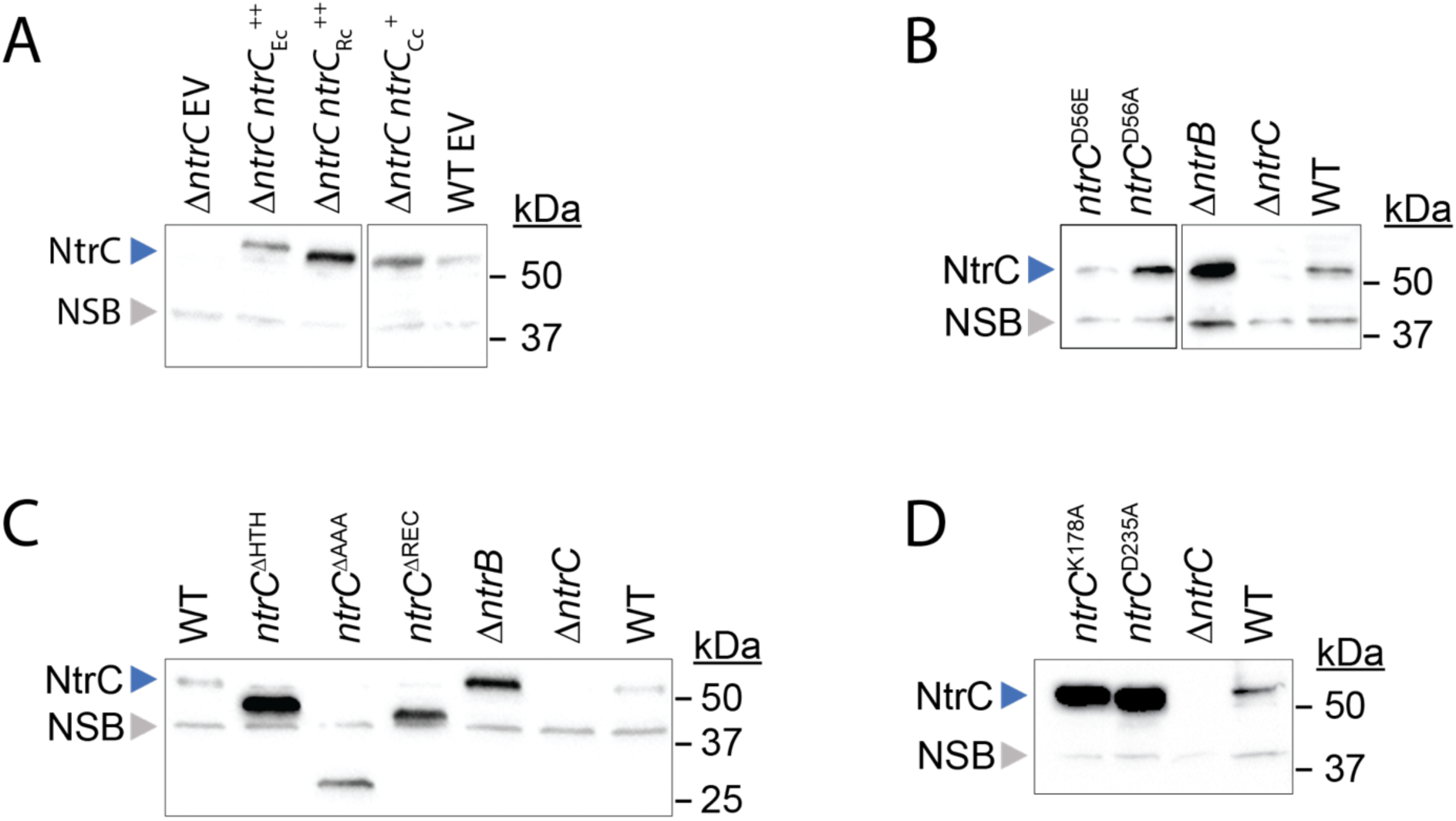
Western blot to assess steady state levels of *Caulobacter* NtrC alleles, and *E. coli* and *R. capsulatus* NtrC expressed in *Caulobacter*. In all panels, wild-type *Caulobacter* NtrC is marked by the blue arrow (∼52 kDa); the non-specific band (NSB) serving as a loading control is marked with a gray arrow. Panel (A) shows an α-NtrC western blot of lysates from *Caulobacter* grown to stationary phase in PYE complex medium supplemented with 0.15% xylose. Displayed are WT and Δ*ntrC* carrying either an empty vector (EV) or expression vectors containing *Caulobacter ntrC* under the control of its native promoter (*ntrC*_Cc_^+^), or *E. coli* or *R. capsulatus ntrC* under the control of a xylose-inducible promoter (*ntrC*_Ec_^++^ or *ntrC*_Rc_^++^). Panels (B-C) present an α-NtrC western blot of lysates WT, Δ*ntrC*, Δ*ntrB*, *ntrC*^D56A^, *ntrC*^D56E^, *ntrC*^ΔREC^ (residues deleted: 17-125), *ntrC*^ΔAAA^ (residues deleted: 159-363), and *ntrC*^ΔHTH^ (residues deleted: 423-462) grown to logarithmic phase in PYE. The predicted molecular weights of NtrC^ΔREC^, NtrC^ΔAAA^, and NtrC^ΔHTH^ are approximately 41 kDa, 30 kDa, and 48 kDa, respectively. Panel (D) shows an α-NtrC western blot of lysates WT, Δ*ntrC*, *ntrC*^K178A^ (Walker A mutant), and *ntrC*^D235A^ (Walker B mutant) grown to stationary phase in PYE.

**Figure S3.**
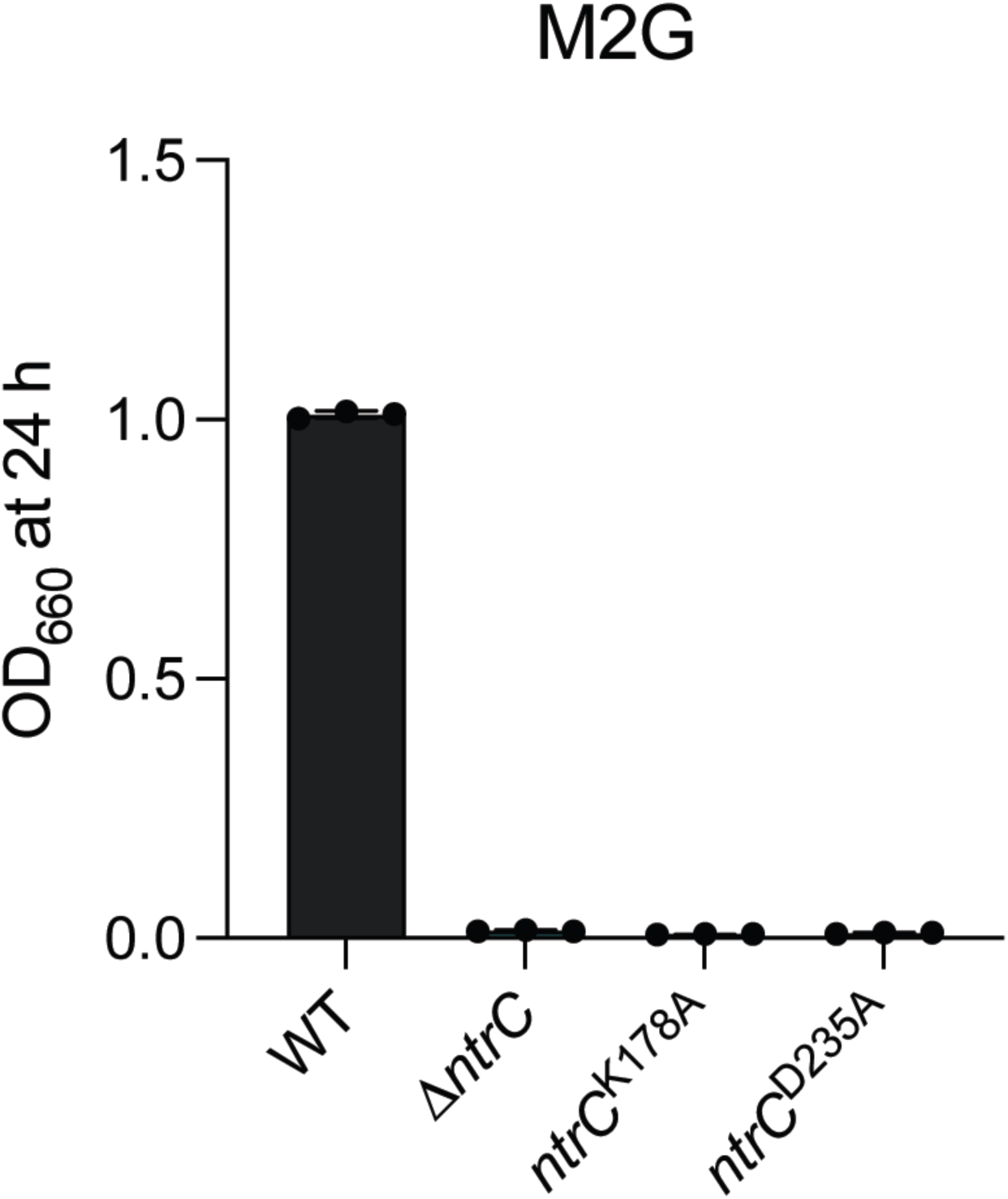
Conserved residues of Walker A and Walker B motifs in the NtrC AAA+ domain are required for growth in defined medium. (A) Terminal OD_660_ of WT, Δ*ntrC*, *ntrC*^K178A^ (Walker A mutant), and *ntrC*^D235A^ (Walker B mutant) after 24 hours (h) of growth in M2G. Data represent mean ± standard deviation of three independent replicates.

**Figure S4.**
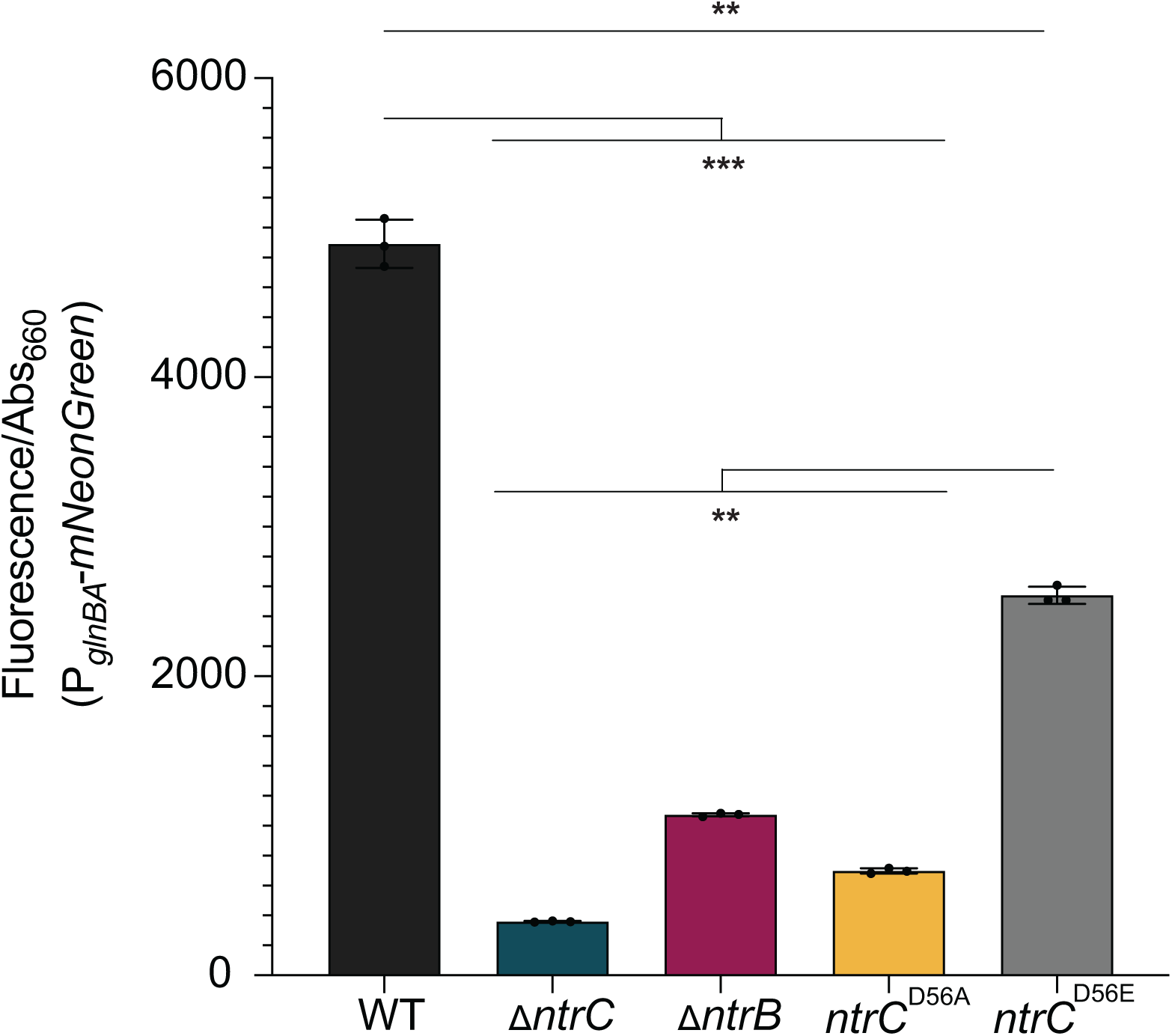
P*_glnBA_* transcription is diminished in *ntrB* and *ntrC* mutant strains. Transcription from the *glnBA* pro-moter (P*_glnBA_*) was measured in WT, Δ*ntrC*, Δ*ntrB*, *ntrC*^D56A^, *ntrC*^D56E^ using a P*_glnBA_*-*mNeonGreen* transcriptional fusion reporter. Strains were grown to stationary phase in PYE. Fluorescence signal was measured and normalized to OD_660_. Data represent mean ± standard deviation of three replicates. Statistical significance was determined by one-way ANOVA followed by Tukey’s multiple comparisons test (*** *P* < .0001, ** *P* ≤ .0008).

**Figure S5.**
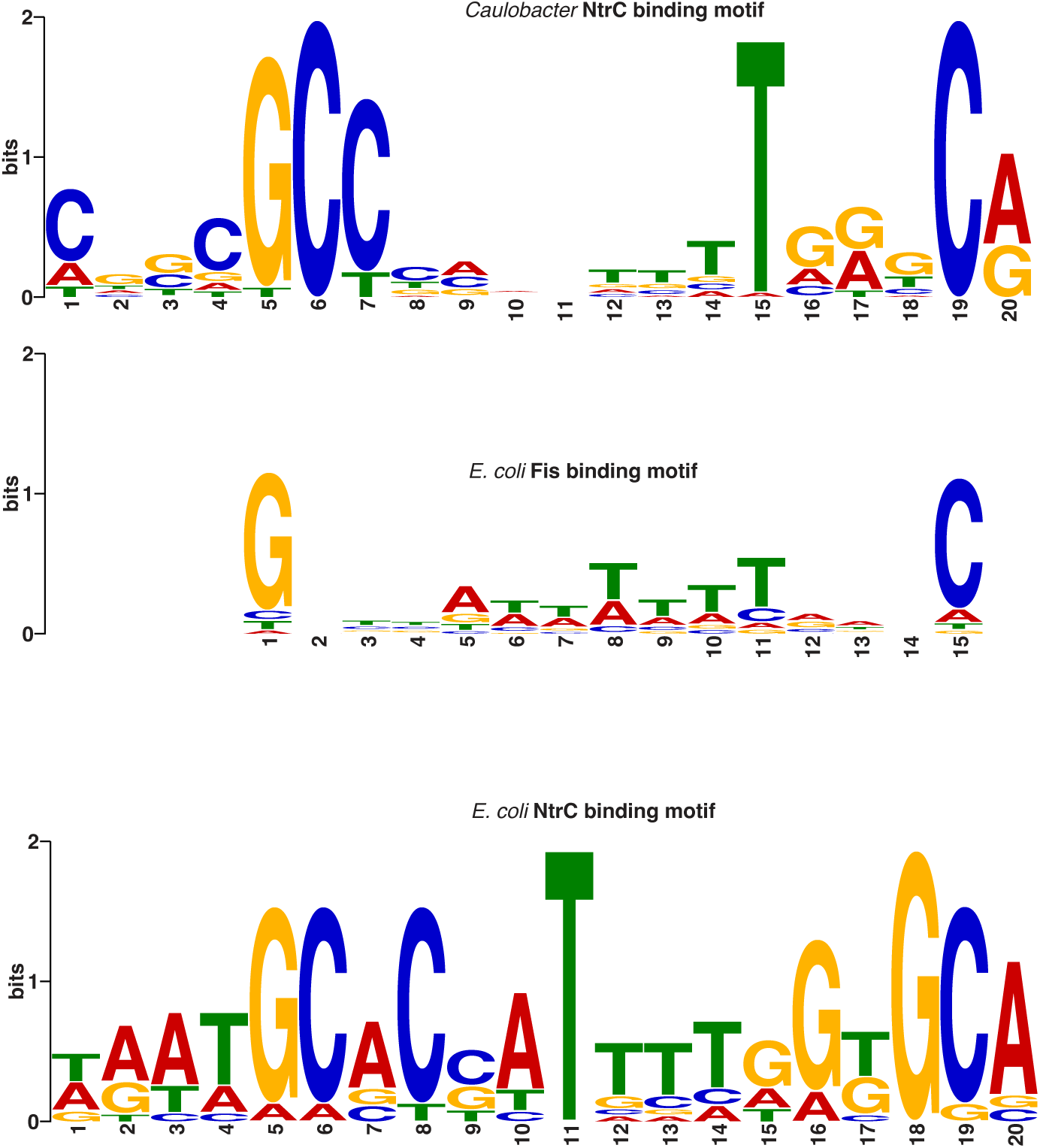
*Caulobacter* NtrC binding motif is similar to *E. coli* Fis and NtrC binding motifs. A search of prokaryotic transcription factor binding motifs in SwissRegulon revealed significant similarity between the *Caulobacter* NtrC binding motif and the Fis and NtrC binding motifs of *E. coli* (*P* < 0.001).

**Figure S6.**
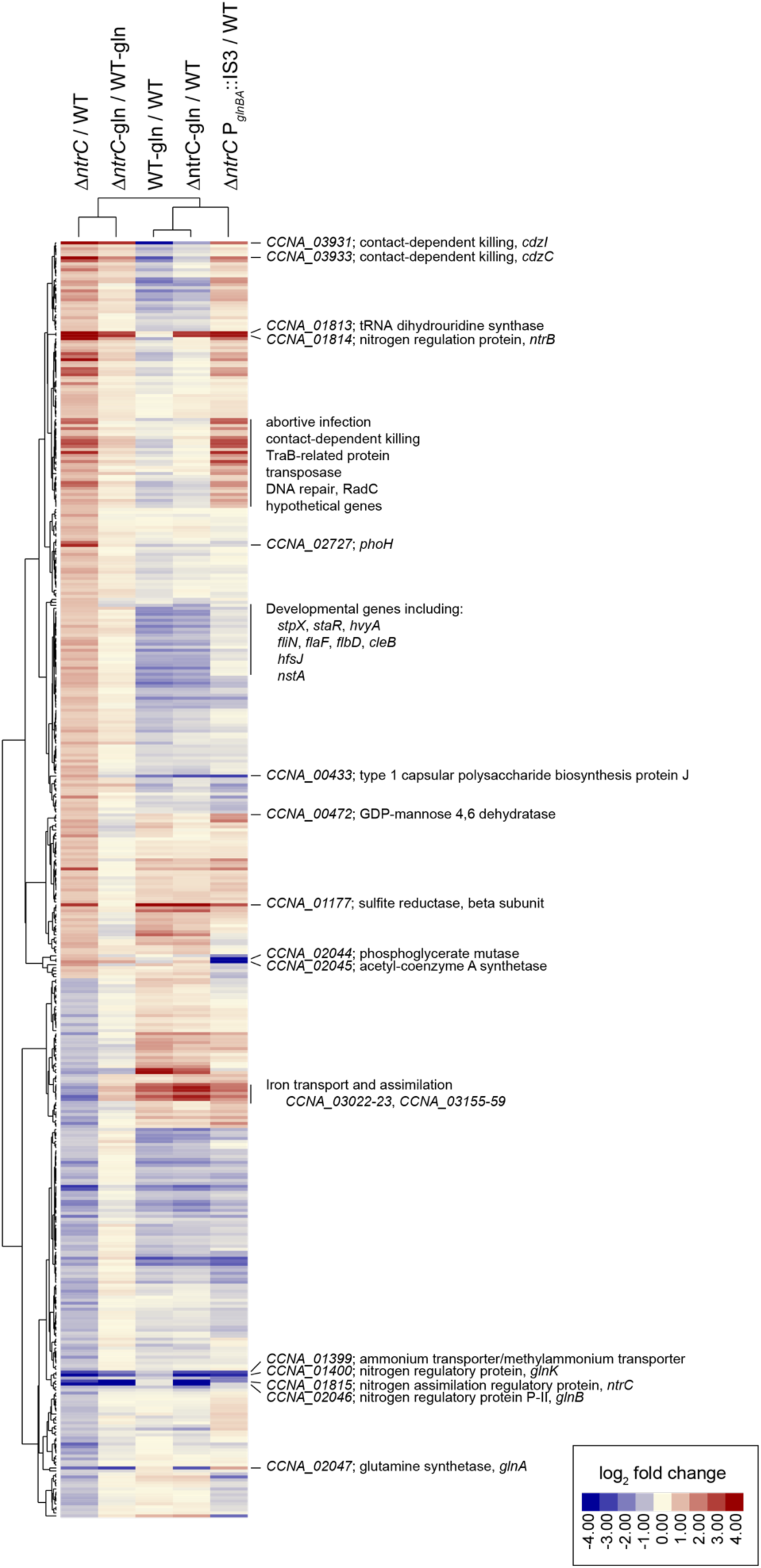
Nitrogen-dependent regulation of the NtrC regulon revealed by RNA-seq analysis. The heat map displays the log_2_ fold change of 473 genes differentially expressed between the Δ*ntrC* mutant and WT. Genes with fold change > 1.5, FDR *P* < 10^-6^, and WT CPM > 10 are included. Each row represents a gene. Each column represents a comparison between strains and/or different media conditions (PYE complex medium and PYE supplemented with 9.3 mM glutamine (gln)). Hierarchical clustering using Cluster 3.0 was applied, employing an uncentered correlation simi-larity metric and average linkage for grouping.

**Figure S7.**
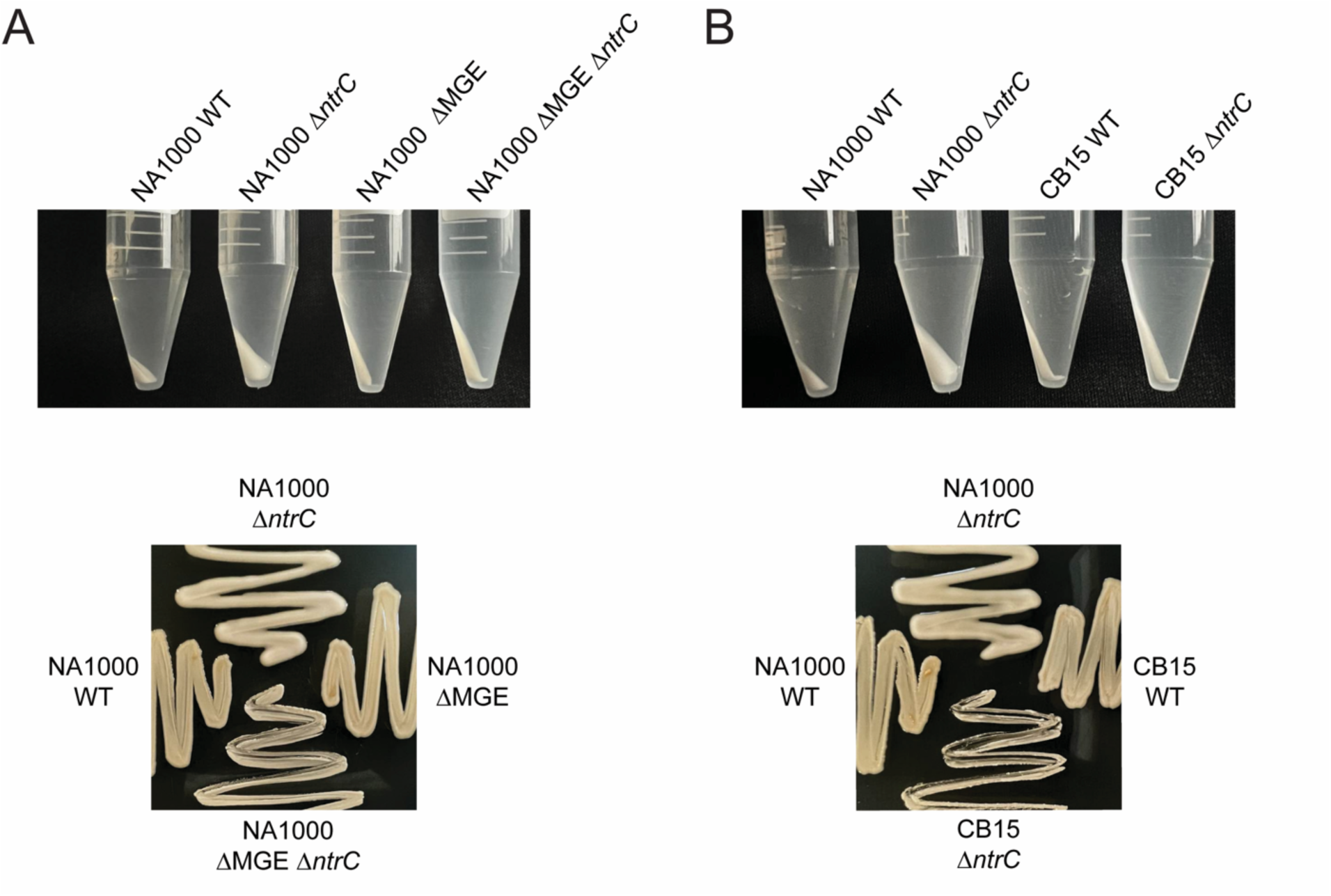
Mucoid phenotype of Δ*ntrC* requires the presence of the 26 kb mobile genetic element (MGE). (A) Top panel: Cell pellets of *Caulobacter crescentus* strain NA1000 WT, NA1000 Δ*ntrC*, NA1000 in which the MGE had spontaneously excised (NA1000 ΔMGE), and NA1000 ΔMGE Δ*ntrC*. Strains were grown overnight in PYE. Overnight cultures were normalized to OD_660_ = 0.5 and cells from 10 ml were centrifuged at 7,197 x g for 3 min at 4°C. Bottom panel: Growth of NA1000 WT, NA1000 Δ*ntrC*, NA1000 ΔMGE Δ*ntrC* on PYE agar supplemented with 3% sucrose. Plates were incubated for 4 days at 30°C. (B) Top panel: Cell pellets of NA1000 WT, NA1000 Δ*ntrC*, CB15 WT, and CB15 Δ*ntrC*. Cell pellets were prepared as described in panel A. Bottom panel: Growth of NA1000 WT, NA1000 Δ*ntrC*, CB15 WT, and CB15 Δ*ntrC* in same conditions described in panel A.

## References

1. Jiang P, Ninfa AJ. 1999. Regulation of autophosphorylation of Escherichia coli nitrogen regulator II by the PII signal transduction protein. J Bacteriol 181:1906–11.

2. Keener J, Kustu S. 1988. Protein kinase and phosphoprotein phosphatase activities of nitrogen regulatory proteins NTRB and NTRC of enteric bacteria: roles of the conserved amino-terminal domain of NTRC. Proc Natl Acad Sci U S A 85:4976–80.

3. Ninfa AJ, Jiang P. 2005. PII signal transduction proteins: sensors of alpha-ketoglutarate that regulate nitrogen metabolism. Curr Opin Microbiol 8:168–73.

4. Schumacher J, Behrends V, Pan Z, Brown DR, Heydenreich F, Lewis MR, Bennett MH, Razzaghi B, Komorowski M, Barahona M, Stumpf MP, Wigneshweraraj S, Bundy JG, Buck M. 2013. Nitrogen and carbon status are integrated at the transcriptional level by the nitrogen regulator NtrC in vivo. mBio 4:e00881–13.

5. von Wiren N, Gazzarrini S, Gojon A, Frommer WB. 2000. The molecular physiology of ammonium uptake and retrieval. Curr Opin Plant Biol 3:254–61.

6. Leigh JA, Dodsworth JA. 2007. Nitrogen regulation in bacteria and archaea. Annu Rev Microbiol 61:349–77.

7. Wilhelm RC. 2018. Following the terrestrial tracks of Cau-lobacter - redefining the ecology of a reputed aquatic oligotroph. ISME J 12:3025–3037.

8. Ronneau S, Petit K, De Bolle X, Hallez R. 2016. Phos-photransferase-dependent accumulation of (p)ppGpp in response to glutamine deprivation in Caulobacter crescentus. Nat Commun 7:11423.

9. Bush M, Dixon R. 2012. The role of bacterial enhancer binding proteins as specialized activators of sigma54-dependent transcription. Microbiol Mol Biol Rev 76:497–529.

10. Aquino P, Honda B, Jaini S, Lyubetskaya A, Hosur K, Chiu JG, Ekladious I, Hu D, Jin L, Sayeg MK, Stettner AI, Wang J, Wong BG, Wong WS, Alexander SL, Ba C, Bensussen SI, Bernstein DB, Braff D, Cha S, Cheng DI, Cho JH, Chou K, Chuang J, Gastler DE, Grasso DJ, Greifenberger JS, Guo C, Hawes AK, Israni DV, Jain SR, Kim J, Lei J, Li H, Li D, Li Q, Mancuso CP, Mao N, Masud SF, Meisel CL, Mi J, Nykyforchyn CS, Park M, Peterson HM, Ramirez AK, Reynolds DS, Rim NG, Saffie JC, Su H, Su WR, et al. 2017. Coordinated regulation of acid resistance in Escherichia coli. BMC Syst Biol 11:1.

11. Brown DR, Barton G, Pan Z, Buck M, Wigneshweraraj S. 2014. Nitrogen stress response and stringent response are coupled in Escherichia coli. Nat Commun 5:4115.

12. Zimmer DP, Soupene E, Lee HL, Wendisch VF, Khodursky AB, Peter BJ, Bender RA, Kustu S. 2000. Nitrogen regulatory protein C-controlled genes of Escherichia coli: scavenging as a defense against nitrogen limitation. Proc Natl Acad Sci U S A 97:14674–9.

13. Dago AE, Wigneshweraraj SR, Buck M, Morett E. 2007. A role for the conserved GAFTGA motif of AAA+ transcription activators in sensing promoter DNA conformation. J Biol Chem 282:1087–97.

14. Cullen PJ, Bowman WC, Kranz RG. 1996. In vitro reconstitution and characterization of the Rhodobacter capsulatus NtrB and NtrC two-component system. J Biol Chem 271:6530–6.

15. Bowman WC, Kranz RG. 1998. A bacterial ATP-dependent, enhancer binding protein that activates the housekeeping RNA polymerase. Genes Dev 12:1884–93.

16. Hsieh ML, Kiel N, Jenkins LMM, Ng WL, Knipling L, Waters CM, Hinton DM. 2022. The Vibrio cholerae master regulator for the activation of biofilm biogenesis genes, VpsR, senses both cyclic di-GMP and phosphate. Nucleic Acids Res 50:4484–4499.

17. Guo MS, Haakonsen DL, Zeng W, Schumacher MA, Laub MT. 2018. A Bacterial Chromosome Structuring Protein Binds Overtwisted DNA to Stimulate Type II Topoisomerases and Enable DNA Replication. Cell 175:583–597 e23.

18. Ricci DP, Melfi MD, Lasker K, Dill DL, McAdams HH, Shapiro L. 2016. Cell cycle progression in Caulobacter requires a nucleoid-associated protein with high AT sequence recognition. Proc Natl Acad Sci U S A 113:E5952–E5961.

19. Fumeaux C, Radhakrishnan SK, Ardissone S, Theraulaz L, Frandi A, Martins D, Nesper J, Abel S, Jenal U, Viollier PH. 2014. Cell cycle transition from S-phase to G1 in Caulobacter is mediated by ancestral virulence regulators. Nat Commun 5:4081.

20. Stewart V. 1994. Regulation of nitrate and nitrite reductase synthesis in enterobacteria. Antonie Van Leeuwenhoek 66:37–45.

21. Cheng AT, Zamorano-Sanchez D, Teschler JK, Wu D, Yildiz FH. 2018. NtrC Adds a New Node to the Complex Regulatory Network of Biofilm Formation and vps Expression in Vibrio cholerae. J Bacteriol 200.

22. Hervas AB, Canosa I, Little R, Dixon R, Santero E. 2009. NtrC-dependent regulatory network for nitrogen assimilation in Pseudomonas putida. J Bacteriol 191:6123–35.

23. Toukdarian A, Kennedy C. 1986. Regulation of nitrogen metabolism in Azotobacter vinelandii: isolation of ntr and glnA genes and construction of ntr mutants. EMBO J 5:399–407.

24. Wardhan H, McPherson MJ, Sastry GR. 1989. Identification, cloning, and sequence analysis of the nitrogen regulation gene ntrC of Agrobacterium tumefaciens C58. Mol Plant Microbe Interact 2:241–8.

25. Capra EJ, Perchuk BS, Skerker JM, Laub MT. 2012. Adaptive mutations that prevent crosstalk enable the expansion of paralogous signaling protein families. Cell 150:222–32.

26. Smith JG, Latiolais JA, Guanga GP, Pennington JD, Silver-smith RE, Bourret RB. 2004. A search for amino acid substitutions that universally activate response regulators. Mol Microbiol 51:887–901.

27. Schumacher J, Zhang X, Jones S, Bordes P, Buck M. 2004. ATP-dependent transcriptional activation by bacterial PspF AAA+protein. J Mol Biol 338:863–75.

28. Rombel I, Peters-Wendisch P, Mesecar A, Thorgeirsson T, Shin YK, Kustu S. 1999. MgATP binding and hydrolysis determinants of NtrC, a bacterial enhancer-binding protein. J Bacteriol 181:4628–38.

29. van Heeswijk WC, Hoving S, Molenaar D, Stegeman B, Kahn D, Westerhoff HV. 1996. An alternative PII protein in the regulation of glutamine synthetase in Escherichia coli. Mol Microbiol 21:133–46.

30. Fiebig A, Castro Rojas CM, Siegal-Gaskins D, Crosson S. 2010. Interaction specificity, toxicity and regulation of a paralogous set of ParE/RelE-family toxin-antitoxin systems. Mol Microbiol 77:236–51.

31. Marks ME, Castro-Rojas CM, Teiling C, Du L, Kapatral V, Walunas TL, Crosson S. 2010. The genetic basis of laboratory adaptation in Caulobacter crescentus. J Bacteriol 192:3678–88.

32. Ardissone S, Fumeaux C, Berge M, Beaussart A, Theraulaz L, Radhakrishnan SK, Dufrene YF, Viollier PH. 2014. Cell cycle constraints on capsulation and bacteriophage susceptibility. Elife 3:e03587.

33. Gora KG, Tsokos CG, Chen YE, Srinivasan BS, Perchuk BS, Laub MT. 2010. A cell-type-specific protein-protein interaction modulates transcriptional activity of a master regulator in Caulobacter crescentus. Mol Cell 39:455–67.

34. Tan MH, Kozdon JB, Shen X, Shapiro L, McAdams HH. 2010. An essential transcription factor, SciP, enhances robustness of Caulobacter cell cycle regulation. Proc Natl Acad Sci U S A 107:18985–90.

35. Collier J, Shapiro L. 2009. Feedback control of DnaA-mediated replication initiation by replisome-associated HdaA protein in Caulobacter. J Bacteriol 191:5706–16.

36. Aakre CD, Phung TN, Huang D, Laub MT. 2013. A bacterial toxin inhibits DNA replication elongation through a direct interaction with the beta sliding clamp. Mol Cell 52:617–28.

37. Garcia-Bayona L, Guo MS, Laub MT. 2017. Contact-dependent killing by Caulobacter crescentus via cell surface-associated, glycine zipper proteins. Elife 6.

38. Anantharaman V, Koonin EV, Aravind L. 2002. Comparative genomics and evolution of proteins involved in RNA metabolism. Nucleic Acids Res 30:1427–64.

39. Andrews ESV, Arcus VL. 2020. PhoH2 proteins couple RNA helicase and RNAse activities. Protein Sci 29:883–892.

40. da Silva Neto JF, Braz VS, Italiani VC, Marques MV. 2009. Fur controls iron homeostasis and oxidative stress defense in the oligotrophic alpha-proteobacterium Caulobacter crescentus. Nucleic Acids Res 37:4812–25.

41. Gonin M, Quardokus EM, O’Donnol D, Maddock J, Brun YV. 2000. Regulation of stalk elongation by phosphate in Caulobacter crescentus. J Bacteriol 182:337–47.

42. de Young KD, Stankeviciute G, Klein EA. 2020. Sugar-Phosphate Metabolism Regulates Stationary-Phase Entry and Stalk Elongation in Caulobacter crescentus. J Bacteriol 202.

43. Ishige T, Krause M, Bott M, Wendisch VF, Sahm H. 2003. The phosphate starvation stimulon of Corynebacterium glutamicum deter-mined by DNA microarray analyses. J Bacteriol 185:4519–29.

44. Kim SK, Makino K, Amemura M, Shinagawa H, Nakata A. 1993. Molecular analysis of the phoH gene, belonging to the phosphate regulon in Escherichia coli. J Bacteriol 175:1316–24.

45. Ravenscroft N, Walker SG, Dutton GG, Smit J. 1991. Identification, isolation, and structural studies of extracellular polysaccharides produced by Caulobacter crescentus. J Bacteriol 173:5677–84.

46. Luria SE, Delbruck M. 1943. Mutations of Bacteria from Virus Sensitivity to Virus Resistance. Genetics 28:491–511.

47. Charlier D, Piette J, Glansdorff N. 1982. IS3 can function as a mobile promoter in E. coli. Nucleic Acids Res 10:5935–48.

48. Boutte CC, Crosson S. 2011. The complex logic of stringent response regulation in Caulobacter crescentus: starvation signalling in an oligotrophic environment. Mol Microbiol 80:695–714.

49. Groisman EA. 2016. Feedback Control of Two-Component Regulatory Systems. Annu Rev Microbiol 70:103–24.

50. Hallgren J, Koonce K, Felletti M, Mortier J, Turco E, Jonas K. 2023. Phosphate starvation decouples cell differentiation from DNA replication control in the dimorphic bacterium *Caulobacter crescentus*. bio-Rxiv 10.1101/2023.07.26.550773.

51. Schlimpert S, Klein EA, Briegel A, Hughes V, Kahnt J, Bolte K, Maier UG, Brun YV, Jensen GJ, Gitai Z, Thanbichler M. 2012. General protein diffusion barriers create compartments within bacterial cells. Cell 151:1270–82.

52. Heinrich K, Leslie DJ, Morlock M, Bertilsson S, Jonas K. 2019. Molecular Basis and Ecological Relevance of Caulobacter Cell Filamentation in Freshwater Habitats. mBio 10.

53. Ireland MM, Karty JA, Quardokus EM, Reilly JP, Brun YV. 2002. Proteomic analysis of the Caulobacter crescentus stalk indicates competence for nutrient uptake. Mol Microbiol 45:1029–41.

54. Wagner JK, Setayeshgar S, Sharon LA, Reilly JP, Brun YV. 2006. A nutrient uptake role for bacterial cell envelope extensions. Proc Natl Acad Sci U S A 103:11772–7.

55. Klein EA, Schlimpert S, Hughes V, Brun YV, Thanbichler M, Gitai Z. 2013. Physiological role of stalk lengthening in Caulobacter crescentus. Commun Integr Biol 6:e24561.

56. Lubin EA, Henry JT, Fiebig A, Crosson S, Laub MT. 2016. Identification of the PhoB Regulon and Role of PhoU in the Phosphate Starvation Response of Caulobacter crescentus. J Bacteriol 198:187–200.

57. Hu P, Brodie EL, Suzuki Y, McAdams HH, Andersen GL. 2005. Whole-genome transcriptional analysis of heavy metal stresses in Caulobacter crescentus. J Bacteriol 187:8437–49.

58. Santos-Beneit F. 2015. The Pho regulon: a huge regulatory network in bacteria. Front Microbiol 6:402.

59. Alford MA, Baghela A, Yeung ATY, Pletzer D, Hancock REW. 2020. NtrBC Regulates Invasiveness and Virulence of Pseudomonas aeruginosa During High-Density Infection. Front Microbiol 11:773.

60. Kim HS, Park SJ, Lee KH. 2009. Role of NtrC-regulated exopolysaccharides in the biofilm formation and pathogenic interaction of Vibrio vulnificus. Mol Microbiol 74:436–53.

61. Liu Y, Lardi M, Pedrioli A, Eberl L, Pessi G. 2017. NtrC-dependent control of exopolysaccharide synthesis and motility in Burkholderia cenocepacia H111. PLoS One 12:e0180362.

62. Herr KL, Carey AM, Heckman TI, Chavez JL, Johnson CN, Harvey E, Gamroth WA, Wulfing BS, Van Kessel RA, Marks ME. 2018. Exopolysaccharide production in Caulobacter crescentus: A resource allocation trade-off between protection and proliferation. PLoS One 13:e0190371.

63. Hottes AK, Shapiro L, McAdams HH. 2005. DnaA coordinates replication initiation and cell cycle transcription in Caulobacter crescentus. Mol Microbiol 58:1340–53.

64. Zweiger G, Shapiro L. 1994. Expression of Caulobacter dnaA as a function of the cell cycle. J Bacteriol 176:401–8.

65. Hsieh ML, Hinton DM, Waters CM. 2018. VpsR and cyclic di-GMP together drive transcription initiation to activate biofilm formation in Vibrio cholerae. Nucleic Acids Res 46:8876–8887.

66. Biondi EG, Skerker JM, Arif M, Prasol MS, Perchuk BS, Laub MT. 2006. A phosphorelay system controls stalk biogenesis during cell cycle progression in Caulobacter crescentus. Mol Microbiol 59:386–401.

67. Ho SN, Hunt HD, Horton RM, Pullen JK, Pease LR. 1989. Site-directed mutagenesis by overlap extension using the polymerase chain reaction. Gene 77:51–9.

68. Thanbichler M, Iniesta AA, Shapiro L. 2007. A comprehensive set of plasmids for vanillate-and xylose-inducible gene expression in Caulobacter crescentus. Nucleic Acids Res 35:e137.

69. McLaughlin M, Hershey DM, Reyes Ruiz LM, Fiebig A, Crosson S. 2022. A cryptic transcription factor regulates Caulobacter adhesin development. PLoS Genet 18:e1010481.

70. Ely B. 1991. Genetics of Caulobacter crescentus. Methods Enzymol 204:372–84.

71. Hmelo LR, Borlee BR, Almblad H, Love ME, Randall TE, Tseng BS, Lin C, Irie Y, Storek KM, Yang JJ, Siehnel RJ, Howell PL, Singh PK, Tolker-Nielsen T, Parsek MR, Schweizer HP, Harrison JJ. 2015. Precision-engineering the Pseudomonas aeruginosa genome with two-step allelic exchange. Nat Protoc 10:1820–41.

72. Pitcher DG, Saunders NA, Owen RJ. 1989. Rapid extraction of bacterial genomic DNA with guanidium thiocyanate. Lett Appl Microbiol 8:151–156.

73. Deatherage DE, Barrick JE. 2014. Identification of mutations in laboratory-evolved microbes from next-generation sequencing data using breseq. Methods Mol Biol 1151:165–88.

74. de Hoon MJ, Imoto S, Nolan J, Miyano S. 2004. Open source clustering software. Bioinformatics 20:1453–4.

75. Saldanha AJ. 2004. Java Treeview--extensible visualization of microarray data. Bioinformatics 20:3246–8.

76. Bharmal MH, Aretakis JR, Schrader JM. 2020. An Improved Caulobacter crescentus Operon Annotation Based on Transcriptome Data. Microbiol Resour Announc 9.

77. Zhou B, Schrader JM, Kalogeraki VS, Abeliuk E, Dinh CB, Pham JQ, Cui ZZ, Dill DL, McAdams HH, Shapiro L. 2015. The global regulatory architecture of transcription during the Caulobacter cell cycle. PLoS Genet 11:e1004831.

78. Zhu LJ, Gazin C, Lawson ND, Pages H, Lin SM, Lapointe DS, Green MR. 2010. ChIPpeakAnno: a Bioconductor package to annotate ChIP-seq and ChIP-chip data. BMC Bioinformatics 11:237.

79. Bailey TL, Johnson J, Grant CE, Noble WS. 2015. The MEME Suite. Nucleic Acids Res 43:W39–49.

80. Hartmann R, van Teeseling MCF, Thanbichler M, Drescher K. 2020. BacStalk: A comprehensive and interactive image analysis software tool for bacterial cell biology. Mol Microbiol 114:140–150.

